# R51Q SNX10 induces osteopetrosis by promoting uncontrolled fusion of monocytes to form giant, non-functional osteoclasts

**DOI:** 10.1101/332551

**Authors:** Maayan Barnea, Merle Stein, Sabina Winograd-Katz, Moran Shalev, Esther Arman, Ori Brenner, Fadi Thalji, Moien Kanaan, Hila Elinav, Polina Stepensky, Benjamin Geiger, Jan Tuckermann, Ari Elson

## Abstract

The molecular mechanisms that regulate fusion of monocytes into functional osteoclasts are virtually unknown. We describe a knock-in mouse model for the R51Q mutation in sorting nexin 10 (SNX10) that exhibits osteopetrosis and related symptoms of patients of autosomal recessive osteopetrosis linked to this mutation. Osteopetrosis arises in homozygous R51Q SNX10 mice due to a unique combination of reduced numbers of osteoclasts that are non-functional. Fusion of mutant monocytes is deregulated and occurs rapidly and continuously to form giant, non-functional osteoclasts. Mutant osteoclasts mature quickly and survive poorly *in vitro*, possibly accounting for their scarcity *in vivo*. These cells also exhibit impaired ruffled borders, which are required for bone resorption, providing an additional basis for the osteopetrotic phenotype. More broadly, we propose that the maximal size of osteoclasts is actively determined by a genetically-regulated, cell-autonomous mechanism that limits precursor cell fusion, and for which SNX10 is required.

## Introduction

Growth and remodeling of bone requires coordinated activities of osteoblasts, which deposit new bone matrix, an osteoclasts (OCLs), which degrade it. OCLs are large, multi-nucleated phagocytic cells that are generated by fusion of monocyte-macrophage precursors in a process driven by the cytokines M-CSF and RANKL. OCLs adhere to mineralized tissue via podosomes, specialized adhesion structures that assemble into a sealing zone at the cell periphery, which confines the bone area that is being degraded. The cells then secrete from a specialized region of their ventral membrane, known as the ruffled border, proteases and protons, which degrade the protein and mineral components of the bone in the resorption cavity (Novack and Teitelbaum, 2008). Despite existence of considerable knowledge concerning OCL formation and function, the mechanisms and molecular players that regulate OCL fusion are mostly unknown.

Proper balance between bone synthesis and degradation is critical for healthy bone homeostasis, and upsetting this balance by excessive or deficient OCL activity leads to skeletal disorders such as osteoporosis or osteopetrosis, respectively (Bruzzaniti and Baron, 2006; Novack and Teitelbaum, 2008). Of particular relevance to this study is autosomal recessive osteopetrosis (ARO; infantile malignant osteopetrosis, IMO), a cluster of genetic disorders that is caused by failure of OCL-mediated bone resorption. ARO is a rare disease, with a global incidence of 1 in 250,000 live births, and is lethal unless treated in a timely manner by hematopoietic stem cell transplantation that enables production of functional osteoclasts (Palagano et al., 2018; Sobacchi et al., 2013). In most ARO cases OCLs are present but are non-functional (“osteoclast-rich ARO”, (Sobacchi et al., 2013)). Roughly two-thirds of these cases are caused by mutations in *TCIRG1* or in CLCN7, which are required for acidification of the resorption cavity. Osteoclast-rich ARO can also arise due to mutations in *OSTM1*, which also participates in acidification, and in *PLEKHM1*, which functions in vesicular trafficking (Del Fattore et al., 2008; Palagano et al., 2018; Sobacchi et al., 2013). Osteoclast-poor ARO, caused by absence of OCLs, has been linked to mutations in *TNFRSF11A* or *TNFSF11*, which encode the TNF receptor superfamily member 11A (RANK) or its ligand, RANKL, respectively. Both molecules are essential for differentiation of OCLs from their monocytic precursor cells (Palagano et al., 2018; Sobacchi et al., 2013).

Homozygosity for the R51Q mutation in Sorting Nexin 10 (SNX10) was shown recently to cause ARO due to inactive OCLs in patients from Palestinian clans (Aker et al., 2012). Subsequent studies have revealed additional mutations in *SNX10*, including nonsense, mis-sense, and splicing defect mutations, that cause ARO in patients of diverse ethnic origins in Europe, North America and Asia. It is currently estimated that mutations in *SNX10* account for 5% of ARO cases globally (Palagano et al., 2018; Pangrazio et al., 2013; Stattin et al., 2017). The specific symptoms of patients carrying distinct *SNX10* mutations vary, but several symptoms are characteristic; these include early age of onset, significantly increased bone mass, anemia, hepatosplenomegaly, and impaired vision and hearing that are caused by gradual closure of skull foramina arising from reduced bone resorption. Additional symptoms may include short stature, failure to thrive, thrombocytopenia, absent or impacted teeth, and mandibular osteomyelitis (Aker et al., 2012; Pangrazio et al., 2013; Stattin et al., 2017).

SNX10 belongs to a family of over 30 related proteins that participate in the regulation of endosome sorting in cells. All SNX proteins contain a phosphatidylinositol-binding (PX) domain that enables them to interact with cellular membrane compartments (Cullen, 2008), but their exact biochemical functions are largely unknown. In addition to its N-terminal PX domain, SNX10 contains a C-terminal domain that is rich in serine residues and in negatively-charged amino acids. The function of this domain is unknown, but it appears to be required for vacuole formation in mammalian cells (Qin et al., 2006). In osteoclasts SNX10 was reported to be associated with the endoplasmic reticulum and to be present in the nucleus (Zhu et al., 2012). The protein was also shown to be required for differentiation of RAW 264.7 cells into osteoclast-like cells and for their resorptive activity *in vitro* (Zhou et al., 2017; Zhu et al., 2012). SNX10 associates with the ATP-dependent vacuolar proton pump of OCLs (V-ATPase) and affects its function (Chen et al., 2012), suggesting that it might be involved in the localization of the V-ATPase to the ruffled border in OCLs (Aker et al., 2012; Palagano et al., 2018; Pangrazio et al., 2013).

It has been reported that osteoclasts prepared *in vitro* from mononuclear leukocytes of R51Q SNX10 patients are scarce, grow poorly, and exhibit significantly reduced resorptive activity when cultured on mineral-coated substrates (Aker et al., 2012). OCLs prepared from patients carrying the distinct Vasterbotten frameshift mutation in *SNX10* (c.212+1G>T, which leads to premature termination of the protein (Stattin et al., 2017)) are two- to five-fold larger than cells derived from healthy individuals, and are non-functional, presumably due to impaired ruffled borders (Stattin et al., 2017). Global knock-down of the orthologous *Snx10* gene in mice resulted in osteopetrosis, due to reduced activity of osteoclasts, and rickets, due to elevated stomach pH and reduced calcium resorption (Ye et al.). Mice in which *Snx10* was specifically knocked-down in OCLs exhibited osteopetrosis without rickets; OCLs from these mice also exhibited impaired ruffled border structure and acidification ability and were non-functional (Ye et al., 2015).

Notably, R51Q and all other known mutations in SNX10 that are associated with ARO are located within the PX domain of the molecule, confirming its importance for osteoclast function. However, since the clinical manifestations of ARO induced by these mutations are variable (Pangrazio et al., 2013) and since the role of SNX10 in OCLs is not well understood, deeper understanding of how distinct mutations affect osteoclast formation, structure, and resorption activity is needed.

In order to examine the specific consequences of the R51Q mutation in SNX10 *in vivo*, we generated R51Q SNX10 knock-in mice. We show here that homozygous R51Q SNX10 mice are severely osteopetrotic due to lack of OCL-mediated bone resorption. This and other aspects of the R51Q SNX10 mouse phenotype closely resemble the human R51Q SNX10 ARO phenotype, confirming the R51Q mutation as a causative factor for ARO. The osteoclast phenotype of these mice is unique since, in contrast to other known mutations in SNX10 and in other proteins that lead to ARO, the R51Q mutation affects both the number and functionality of these cells. Results presented here suggest that the scarcity of OCLs in homozygous R51Q SNX10 mice is caused by deregulated fusion that results in formation of giant, unstable OCLs. The inability of these cells to resorb bone is attributable to both the altered morphology of the mature OCLs and to their lack of ruffled borders, which are required for bone resorption. More broadly, the large size of these mutant cells indicates that OCL fusion is actively regulated by cellular mechanisms that limit fusion of precursor cells, which require SNX10 and that are disrupted by its R51Q mutation.

## Results

### Impaired growth and development in mice homozygous for the R51Q SNX10 mutation

In order to generate a pre-clinical mouse model for ARO caused by the R51Q SNX10 mutation, we inserted this mutation into exon 4 of the endogenous *Snx10* gene using CRISPR technology (Figure 1 A). Two separate R51Q SNX10 mutant lines of mice were generated from independent founder mice (lines 43 and 87), and both exhibited similar phenotypes. Most of the studies below were performed in mice from line 43, with key results verified also in line 87 as indicated.

**Figure 1:**
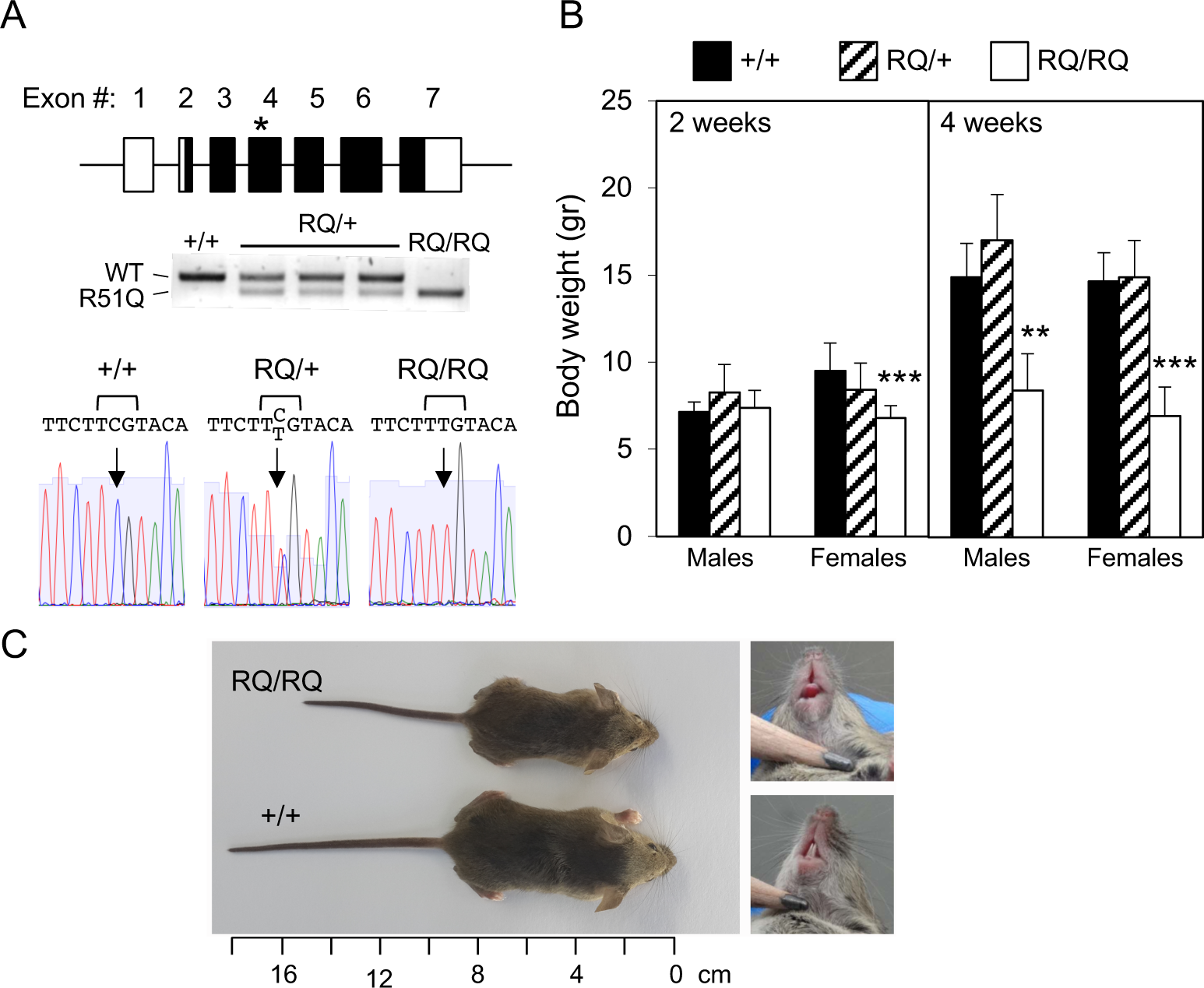
R51Q knock-in mice are growth-retarded. **(A)** Top: Scheme of the *Snx10* gene structure and location of the R51Q mutation (asterisk). Black and white boxes: Coding and non-coding exon segments, respectively; not drawn to scale. Middle: Genotyping of mice by tail biopsy PCR (+/+: wild type; RQ/+: heterozygous; RQ/RQ: homozygous mutant). Bottom: (Anti-sense) sequence from tail biopsy DNA documenting the mutation. The mutant nucleotide is indicated by an arrow. **(B)** Body weights of mice at 2 and 4 weeks of age. Shown are means±SD, N=7-22 mice/bar. **: P<0.0071, ***: P<0.0001 by one-way ANOVA with Tukey’s multiple comparisons test. Data is from lines 43 and 87. **(C)** RQ/RQ mice are shorter than matched +/+ controls and lack teeth. Male mice aged 5 weeks are shown. See also Figure 1-figure supplements 1, 2.

Mating of heterozygous R51Q SNX10 mice within each line yielded wild-type (+/+), heterozygous (RQ/+), and homozygous (RQ/RQ) mutant mice in near-Mendelian ratios, indicating that R51Q SNX10 does not significantly affect prenatal and early post-natal survival. Homozygous RQ/RQ mice appeared grossly normal at birth and during the initial post-natal period (Figure 1B). However, by the age of 4 weeks, significant lags in weight and body length became evident in RQ/RQ mice of both sexes (Figures 1B, 1C; Figure 1 – figure supplement 1). Internal organs, such as the liver, spleen, and stomach, were proportionally smaller in RQ/RQ mice (Figure 1 – figure supplement 2, Table 1). All RQ/RQ mice lacked teeth (Figure 1C), which rendered them dependent on mothers’ milk or crushed lab chow for survival. Histological studies revealed that molars were present but failed to erupt, and osteopetrosis was detected in most skull bones, including the maxillae and mandibles (Figures 2A, 2B). Some RQ/RQ mice developed mandibular osteomyelitis, which involved the osteopetrotic bone and surrounding soft tissues (Figure 2B). RQ/RQ mice were grown routinely up to the age of 6-8 weeks in the presence of lactating females, and anecdotal experience indicated that some can survive longer. Absence of teeth is characteristic of mice suffering from severely reduced OCL-mediated bone resorption (e.g., (Soriano et al., 1991)), suggesting that the RQ/RQ phenotype is associated with OCL dysfunction. Heterozygous RQ/+ mice were physically indistinguishable from wild-type mice by all of the parameters described above.

**Table 1.** Stomach weight, gastric pH, and circulating levels of calcium, parathyroid hormone (PTH) and Vitamin D3 in R51Q SNX10 mice. Data shown is mean ± SE. Mice analyzed were 3 weeks old and included males and females from both lines 43 and 87. All mice were analyzed prior to weaning and had access to mother’s milk throughout. NS: not statistically significant. Statistical significance was determined by two-tailed Student’s t-test.

| Parameters | N | Genotype | Values |
| --- | --- | --- | --- |
| Stomach weight (mg) | 13 | +/+ | 143.1±6.6 |
|  | 8 | *RQ/RQ* | 121.9±9.1 (NS, P=0.075) |
| Body weight (gr) | 13 | +/+ | 11.6±0.4 |
|  | 8 | *RQ/RQ* | 8.7±0.7 (P=0.0022) |
| Stomach weight/body weight (mg/gr) | 13 | +/+ | 12.7±0.8 |
|  | 8 | *RQ/RQ* | 14.7± 1.1 (NS) |
| Gastric pH | 13 | +/+ | 3.10±0.06 |
|  | 8 | *RQ/RQ* | 3.21±0.11 (NS) |
| Serum calcium (mmol/l) | 10 | +/+ | 2.83±0.16 |
|  | 10 | *RQ/RQ* | 2.89±0.12 (NS) |
| Plasma PTH (pg/ml) | 32 | +/+ | 34.77±3.73 |
|  | 14 | *RQ/RQ* | 29.00±9.23 (NS) |
| Plasma 25(OH) Vitamin D (ng/ml) | 32 | +/+ | 14.25±2.43 |
|  | 14 | *RQ/RQ* | 6.97±2.24 (P=0.021) |

**Figure 2:**
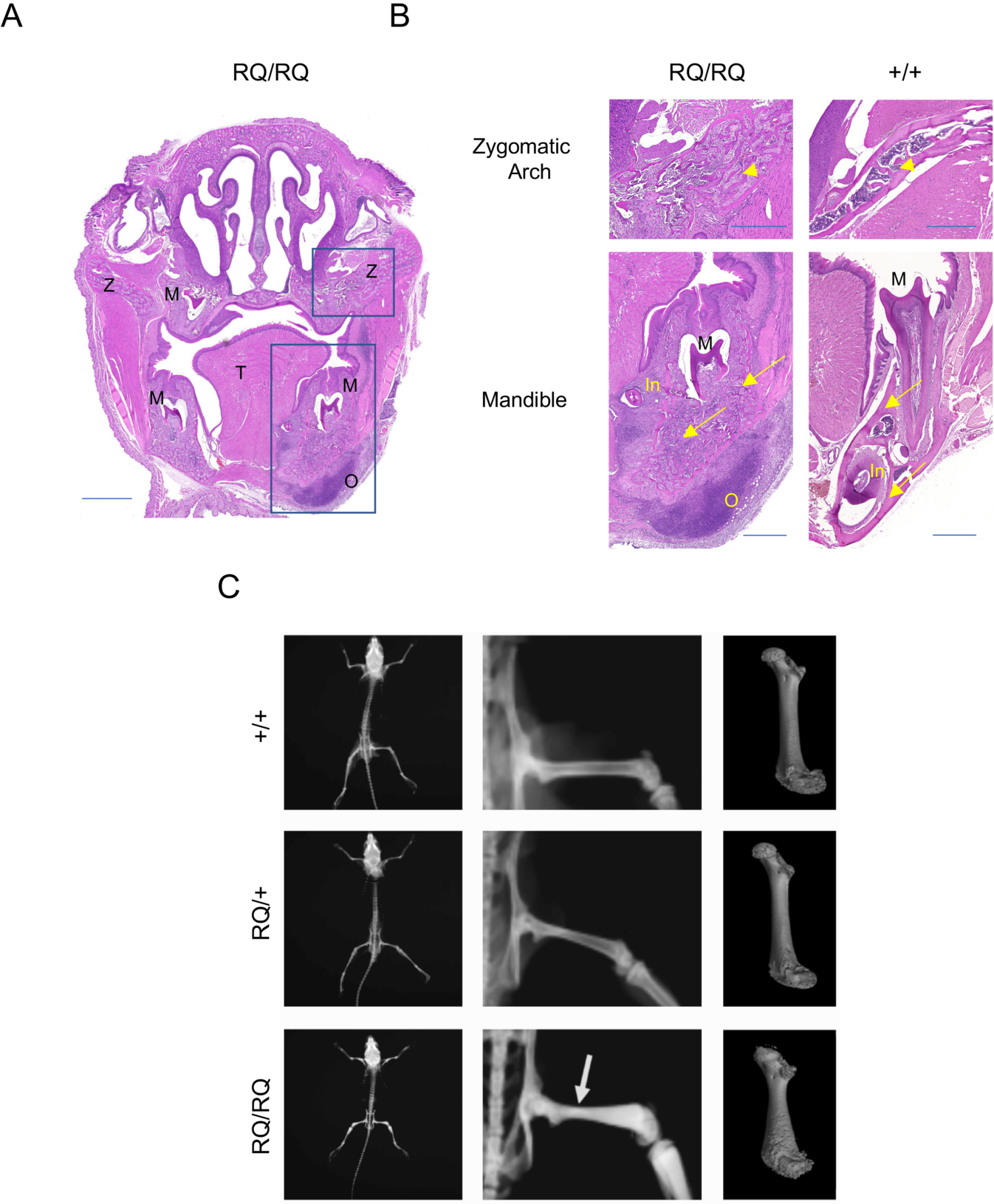
Effect of Snx10 R51Q on bone shape and mineralization. **(A)** Coronal section through RQ/RQ mouse head showing osteopetrosis of the zygomatic arch (upper box) and penetrating mandibular osteomyelitis (lower box). M: Impacted molars; O: Osteomyelitis and cellulitis; T: Tongue; Z: Zygomatic arch. Scale bar: 1,000 μm. **(B)** Enlarged images of boxed regions in Panel A, compared with +/+ controls. Top: Osteopetrosis of the Zygomatic arch. Arrowheads mark examples of osteopetrotic bone (RQ/RQ) vs. normal bone (+/+). Bottom: Mandibular osteomyelitis and cellulitis. In: Incisor (ectopic in RQ/RQ); M: Molar (impacted in RQ/RQ). O: Osteomyelitic region. Arrows mark osteopetrotic and inflamed mandibular bone and adjacent soft tissue (RQ/RQ) vs. normal mandible (+/+). Scale bars: 500 μm. Images shown are representative of 4 RQ/RQ and 4 +/+ mice, age 5-8 weeks. **(C)** Mouse whole skeleton X-ray (left, middle) and 3D femur reconstruction by µCT (right), showing abnormal “Erlenmeyer flask” shape and higher radiodensity (arrow) with severely reduced marrow cavity volume in RQ/RQ mice. Mice shown are females aged 5 weeks, and are representative of 8+/+, 8 RQ/+ and 9 RQ/RQ mice of both sexes.

### Homozygous R51Q SNX10 mice exhibit massive osteopetrosis with mildly reduced bone mineral density

X-ray analysis revealed markedly increased density in bones from RQ/RQ mice relative to RQ/+ and wild-type mice (Figure 2C). In particular, femurs of RQ/RQ mice possessed the ARO “Erlenmeyer flask” shape (Figure 2C;(Faden et al., 2009)) that is also found in human osteopetrotic patients (Sobacchi et al., 2013; Stattin et al., 2017). Micro-CT analysis revealed that femurs from RQ/RQ mice were shorter, deformed, and displayed a somewhat “pockmarked” appearance at the cortical bone surface; the cortical mineral density of these bones was reduced (Figures 2C-E; Table 2). The femoral bone marrow cavities of RQ/RQ mice were almost completely replaced by compact bone matrix (Figure 3A). In line with these observations, the bone volume, trabecular thickness and number of trabeculae per unit of bone volume were significantly increased in these samples, rendering their inter-trabecular separation less apparent (Figure 3B and Table 3). Similar findings were also obtained in vertebrae (Figure 3C and Table 3). Histological analyses of the proximal tibias and vertebras of RQ/RQ mice confirmed massively increased amounts of trabecular bone (Figures 3D, 3E, Figure 3 – figure supplement 1). Significant regions of homozygous trabecular bone stained less strongly with hematoxylin/eosin (Figure 3E, Figure 3 – figure supplement 1), most likely due to the presence of residual unresorbed cartilage. RQ/RQ mice of both sexes were affected to similar extents, and none of the above findings were observed in RQ/+ or +/+ mice.

**Table 2.** Cortical mineral density is reduced in RQ/RQ tibias. Parameters are shown as tissue mineral density in g/cm^3^, normalized to 0.25 and 0.75 g/cm^3^ phantoms. *: P<0.004 relative to +/+ or RQ/+ mice, respectively, by one-way ANOVA with Tukey’s multiple comparisons test. Data are shown as mean ± SD. N=3-5 5-week old mice as indicated.

| Sex | N | Genotype | Tissue Mineral Density<br>(CaHA, g/cm<sup>3</sup>) |
| --- | --- | --- | --- |
| Male | 3 | +/+ | 0.9003± 0.0377 |
|  | 4 | *RQ*/+ | 0.8931± 0.0211 |
|  | 5 | *RQ/RQ* | 0.7690± 0.0223 * |
| Female | 5 | +/+ | 0.9064± 0.0299 |
|  | 4 | *RQ*/+ | 0.9095± 0.0376 |
|  | 4 | *RQ/RQ* | 0.7764± 0.0276 * |

**Table 3.**
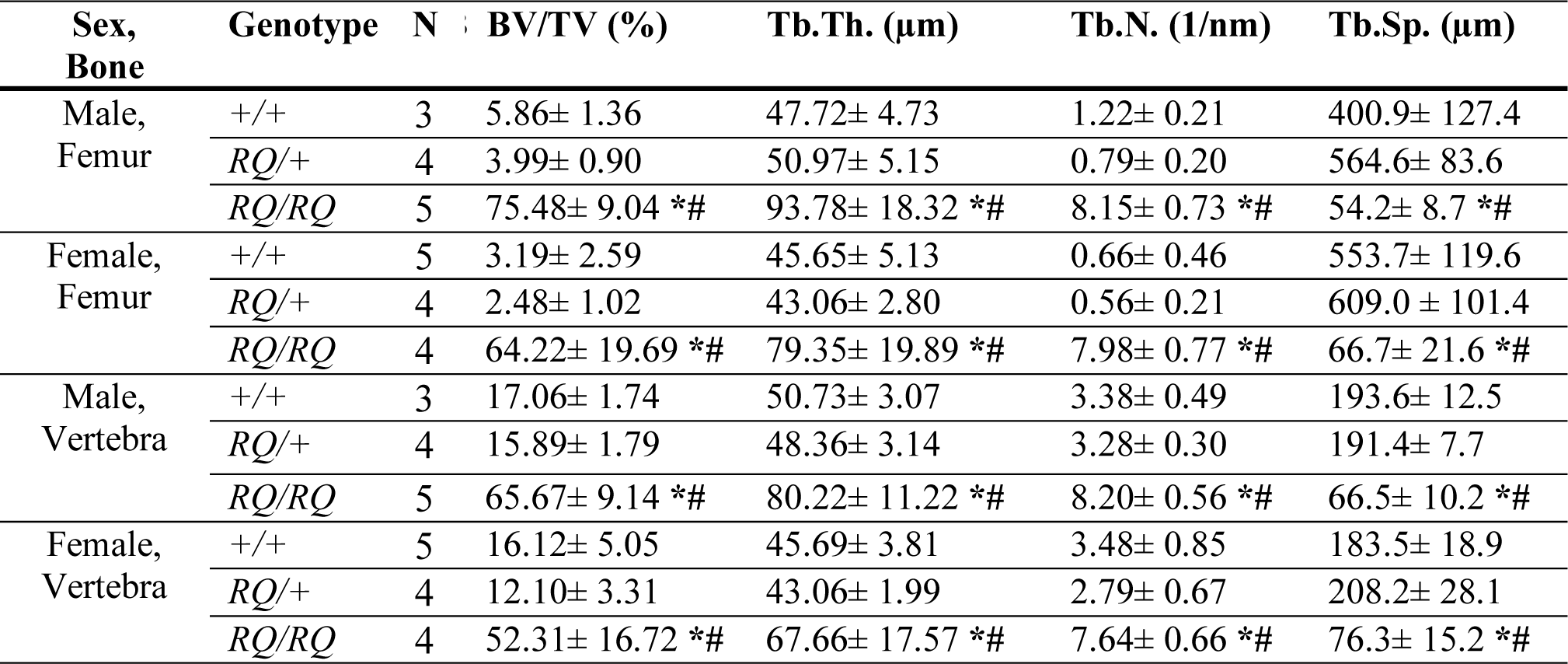
Cancellous bone parameters measured by micro CT reveal osteopetrosis in RQ/RQ mice. Parameters are shown as bone volume per tissue volume (BV/TV), trabecular thickness (Tb.Th.), trabecular number (Tb.N.), and trabecular separation (Tb.Sp.). *, #: P<0.05 relative to +/+, RQ/+ mice, respectively, by the Mann-Whitney-U test. Data are shown as mean ± SD. N=3-5 5-week old mice as indicated.

| Sex,<br>Bone | Genotype | N | BV/TV (%) | Tb.Th. (μm) | Tb.N. (1/nm) | Tb.Sp. (μm) |
| --- | --- | --- | --- | --- | --- | --- |
| Male,<br>Femur | +/+ | 3 | 5.86± 1.36 | 47.72± 4.73 | 1.22± 0.21 | 400.9± 127.4 |
|  | *RQ*/+ | 4 | 3.99± 0.90 | 50.97± 5.15 | 0.79± 0.20 | 564.6± 83.6 |
|  | *RQ/RQ* | 5 | 75.48± 9.04 *# | 93.78± 18.32 *# | 8.15± 0.73 *# | 54.2± 8.7 *# |
| Female,<br>Femur | +/+ | 5 | 3.19± 2.59 | 45.65± 5.13 | 0.66± 0.46 | 553.7± 119.6 |
|  | *RQ*/+ | 4 | 2.48± 1.02 | 43.06± 2.80 | 0.56± 0.21 | 609.0± 101.4 |
|  | *RQ/RQ* | 4 | 64.22± 19.69 *# | 79.35± 19.89 *# | 7.98± 0.77 *# | 66.7± 21.6 *# |
| Male,<br>Vertebra | +/+ | 3 | 17.06± 1.74 | 50.73± 3.07 | 3.38± 0.49 | 193.6± 12.5 |
|  | *RQ*/+ | 4 | 15.89± 1.79 | 48.36± 3.14 | 3.28± 0.30 | 191.4± 7.7 |
|  | *RQ/RQ* | 5 | 65.67± 9.14 *# | 80.22± 11.22 *# | 8.20± 0.56 *# | 66.5± 10.2 *# |
| Female,<br>Vertebra | +/+ | 5 | 16.12± 5.05 | 45.69± 3.81 | 3.48± 0.85 | 183.5± 18.9 |
|  | *RQ*/+ | 4 | 12.10± 3.31 | 43.06± 1.99 | 2.79± 0.67 | 208.2± 28.1 |
|  | *RQ/RQ* | 4 | 52.31± 16.72 *# | 67.66± 17.57 *# | 7.64± 0.66 *# | 76.3± 15.2 *# |

**Figure 3:**
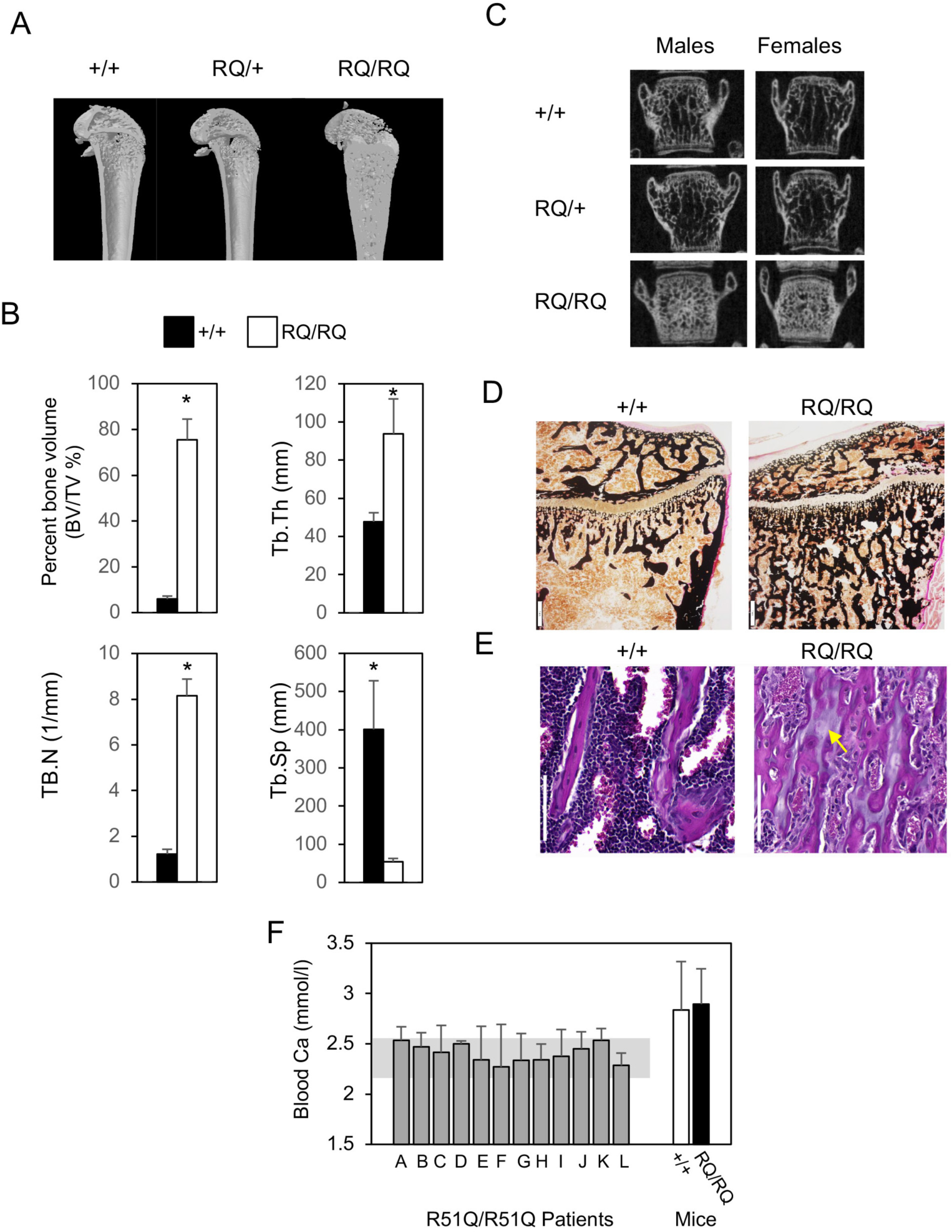
Structure of bones from homozygous R51Q SNX10 mice. Frontal plane section of tibiae from +/+, RQ/+, and RQ/RQ mice by micro-CT. Note nearly complete absence of bone marrow cavity in the homozygous sample. Images are representative of 8+/+, 8 RQ/+ and 8 RQ/RQ mice of both sexes. Key histomorphometric parameters of cancellous bone in femora of male mice aged 5 weeks. Data are mean±SD; N=3-5 mice/bar. *: P<0.001 by Mann-Whitney U test. BV/TV: Bone volume relative to total volume measured. Tb.Th: Trabecular thickness. Tb.N: Number of trabeculae. Tb.Sp: Trabecular separation. See Table 3 for complete data. **(C)** µCT examination of vertebrae, showing increased bone mass and radiodensity in RQ/RQ samples. See Table 3 for complete data. Images represent 8+/+, 8 RQ/+ and 8 RQ/RQ mice of both sexes. **(D)** von Kossa-stained section of tibias from +/+ and RQ/RQ mice. Scale bar: 40 μm. **(E)** Similar to D stained with hematoxylin-eosin, showing magnified view of trabecular bone from a region distal to the growth plate. Arrow denotes unresorbed cartilage. Scale bar: 100μm. **(F)** Serum calcium levels in 12 individual human SNX10 R51Q patients (each bar represents means±SD of repeated measurements of an individual patient over a period of several months). Gray background band shows the healthy laboratory reference range, indicating that all patients exhibit normal values. Right: Snx10 RQ/RQ mice exhibit normal serum calcium levels (N=9-10 mice per genotype, aged 3 weeks). See also Figure 3-figure supplements 1, 2.

SNX10 knockdown (SNX10-KD) mice suffer from a combination of massive osteopetrosis and, paradoxically, rickets due to reduced calcium resorption (Ye et al., 2015). While RQ/RQ mice exhibited massive osteopetrosis and their bone mineral density was somewhat reduced (Table 2), their gastric pH, stomach appearance, and circulating levels of calcium and PTH were normal (Table 1 and Figure 3 – figure supplement 2), collectively indicating that dietary calcium resorption is normal. RQ/RQ mice exhibited reduced levels of circulating 25(OH)-vitamin D (Table 1), which explains their reduced bone mineral density; the reasons for reduced vitamin D levels are unknown. Of note, patients of ARO that is caused by the R51Q SNX10 mutation exhibit normal values of circulating calcium (Figure 3F), and there are no reports of them suffering from rickets. We conclude that RQ/RQ mice suffer from massive and widespread osteopetrosis that is similar to the major clinical symptoms of ARO caused by R51Q SNX10 in humans.

### OCL numbers are decreased in RQ/RQ mice

OCL numbers in bones of RQ/RQ mice were significantly reduced (Figures 4A and 4B, and Table 4). Homozygous bone samples contained normal numbers of osteocytes (Figure 4C, Table 4) but, unexpectedly, reduced numbers of osteoblasts (OBs; Figure 4D; Table 4). However, bone formation rates were normal in RQ/RQ mice (Figure 4E), indicating that the phenotype of these mice is not due to a major bone formation defect. In agreement, OBs derived from calvarias of +/+, RQ/+, and RQ/RQ mice exhibited similar mineralization *in vitro* (Figure 4 – figure supplement 1), and plasma levels of N-terminal type-I procollagen (P1NP), a marker of bone formation *in vivo*, were not significantly distinct in RQ/RQ mice vs. controls (Figure 4F).

**Table 4.**
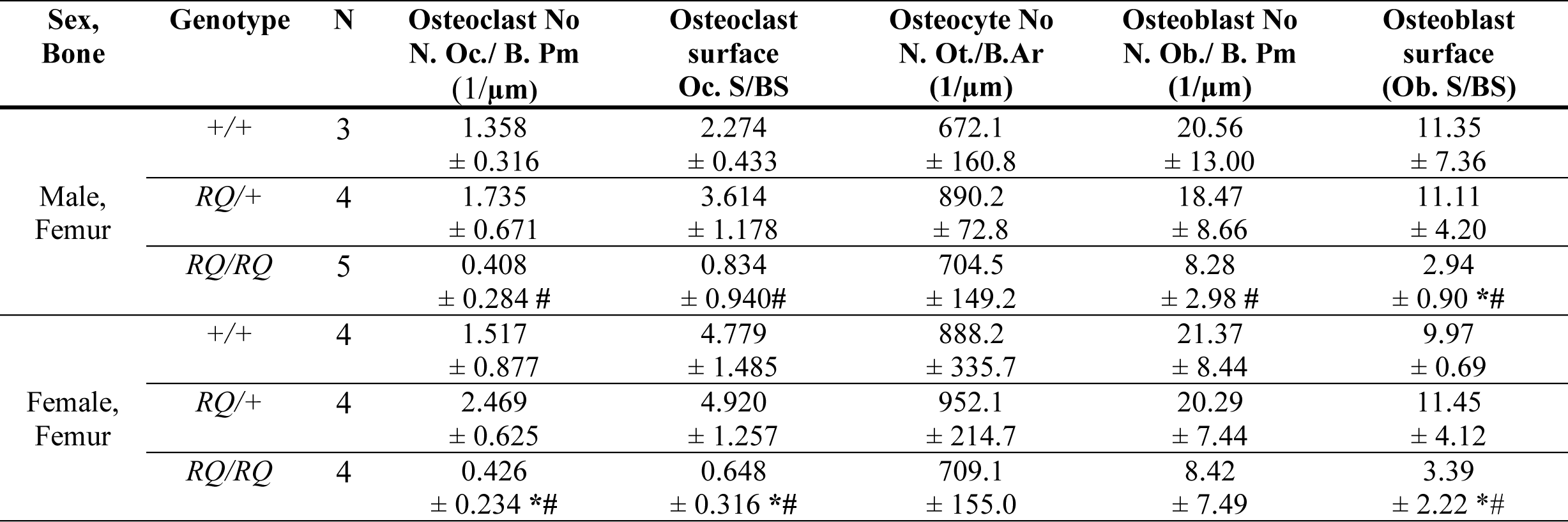
Bone histomorphometry reveals osteoclast and osteoblast abnormalities in RQ/RQ mice. Parameters are shown as Osteoclast number per bone perimeter (N. Oc./ B. Pm), osteoclast surface per bone surface (Oc. S/BS), osteocyte number per bone area (N. Ot./ B. Ar), osteoblast number per bone perimeter (N. Ob./ B. Pm), osteoblast surface per bone surface (Ob. S/BS). *, #: P<0.05 relative to +/+ or RQ/+ mice, respectively, by the Mann-Whitney-U test. Data are shown as mean ± SD. N=3-5 5-week old mice as indicated

| Sex,<br>Bone | Genotype | N | Osteoclast No<br>N. Oc./ B. Pm<br>(1/μm) | Osteoclast<br>surface<br>Oc. S/BS | Osteocyte No<br>N. Ot./B.Ar<br>(1/μm) | Osteoblast No<br>N. Ob./ B. Pm<br>(1/μm) | Osteoblast<br>surface<br>(Ob. S/BS) |
| --- | --- | --- | --- | --- | --- | --- | --- |
| Male,<br>Femur | +/+ | 3 | 1.358<br>± 0.316 | 2.274<br>± 0.433 | 672.1<br>± 160.8 | 20.56<br>± 13.00 | 11.35<br>± 7.36 |
|  | *RQ*/+ | 4 | 1.735<br>± 0.671 | 3.614<br>± 1.178 | 890.2<br>± 72.8 | 18.47<br>± 8.66 | 11.11<br>± 4.20 |
|  | *RQ*/*RQ* | 5 | 0.408<br>± 0.284 # | 0.834<br>± 0.940# | 704.5<br>± 149.2 | 8.28<br>± 2.98 # | 2.94<br>± 0.90 *# |
| Female,<br>Femur | +/+ | 4 | 1.517<br>± 0.877 | 4.779<br>± 1.485 | 888.2<br>± 335.7 | 21.37<br>± 8.44 | 9.97<br>± 0.69 |
|  | *RQ*/+ | 4 | 2.469<br>± 0.625 | 4.920<br>± 1.257 | 952.1<br>± 214.7 | 20.29<br>± 7.44 | 11.45<br>± 4.12 |
|  | *RQ*/*RQ* | 4 | 0.426<br>± 0.234 *# | 0.648<br>± 0.316 *# | 709.1<br>± 155.0 | 8.42<br>± 7.49 | 3.39<br>± 2.22 *# |

**Figure 4:**
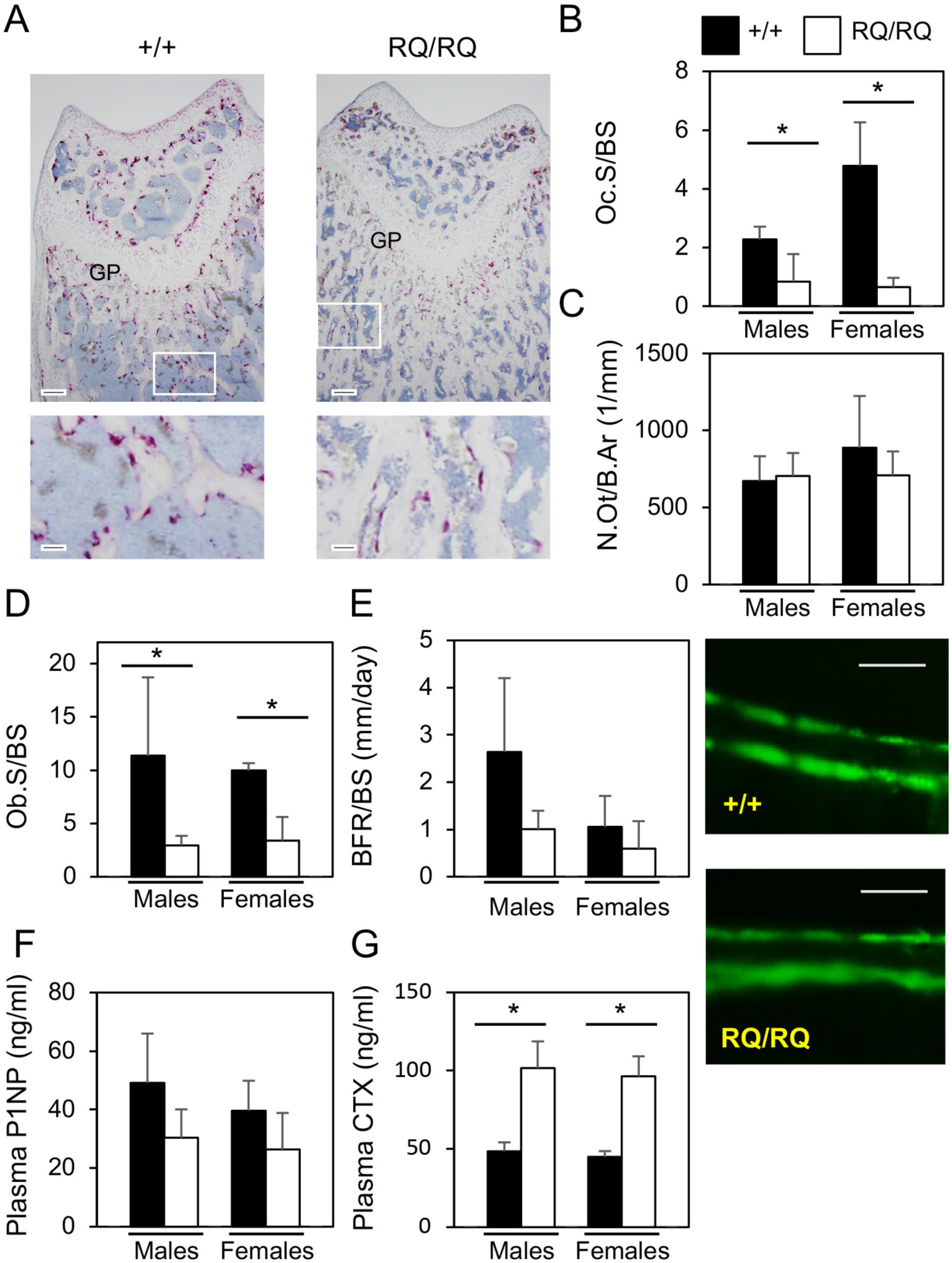
Bone formation and resorption in R51Q SNX10 mice. **(A)** Femur sections from +/+ and homozygous RQ/RQ male mice stained for tartarate-resistant acid phosphatase (TRAP, purple, stains OCLs), and counterstained with hematoxylin. Scale bar: 100 μm. Bottom: Magnified view of boxed regions in top panels. Scale bar: 20 μm. Note reduced OCL numbers in mutant samples. Images are representative of 8 +/+ mice and 9 RQ/RQ mice. **(B)** Histomorphometric quantification of bone surface area in contact with OCLs (Oc.S/BS). *: P≤0.05 by Mann-Whitney U test. (**C**) Histomorphometric quantification of the number of osteocytes per unit bone area (N.Ot/B.Ar). (**D**) Similar to B, for osteoblasts. *: P≤0.05, by Mann-Whitney U test. (**E**) Left: Cortical bone formation rate. Right: incorporation of calcein into bone following two calcein injections 8 days apart. Scale bar: 10 μm. (**F**) Plasma concentrations of N-terminal type-I procollagen (P1NP). **(G)** Plasma concentrations of collagen type-I cross-linked C-telopeptides (CTX). N=11-25 mice age 21-26 days per bar. P≤ 0.011 by Student’s t-test within each sex. N=3-5 mice aged 5 weeks per bar in panels B, C, D, F. All data shown are mean±SD. See also Table 4 and Figure 4 – figure supplement 1.

RQ/RQ mice displayed elevated levels of plasma collagen type-I cross-linked C-telopeptides (CTX) (Figure 4G), which is used as a clinical indicator of OCL-mediated bone resorption *in vivo*. Yet, RQ/RQ mice exhibit a massive increase in bone mass, and their OCLs cannot resorb bone (see below). CTX levels can be affected also by other parameters, such as increased stability of CTX peptides or their reduced clearance rates (Vasikaran et al., 2011). Accordingly, high CTX levels *in vivo* despite reduced OCL activity have been documented in various model systems that exhibit osteopetrosis, including mice lacking CLCN-7 (Neutzsky-Wulff et al., 2008) and in SNX10-KD mice (Ye et al., 2015)).

### RQ/RQ osteoclasts are gigantic

In humans, the cause of ARO is aberrant bone resorption. Absence of teeth in RQ/RQ mice (Figure 1C), reduced OCL numbers in femurs (Figures 4A and 4B), and presence of massive osteopetrosis despite apparently normal bone formation rates (Figure 4E) all suggest that OCL activity is reduced in these mice *in vivo*. In order to examine RQ/RQ OCLs we produced OCLs *in vitro* by culturing primary splenocytes in the presence of M-CSF and RANKL. Cells from RQ/RQ mice adhered to bone and grew rapidly, fusing to form massive multinuclear cells that were dramatically larger than multinucleated cells from +/+ or from RQ/+ mice (Figures 5A, 5B, and 5C). These large RQ/RQ OCLs stained positively for TRAP, but staining was less intense and more diffuse than that seen in +/+ or RQ/+ cells (Figure 5A). RQ/RQ cells seeded on bone grew, and then fused and died before their matched +/+ and RQ/+ counterparts (Figure 5A). Growing +/+ and RQ/+ cells for longer periods of time beyond those of Figure 5A did not produce OCLs larger than those seen in the figure. Each RQ/RQ OCL exhibited a single sealing zone structure that demarcated the entire periphery of the cell, in proportion with its size, in contrast to the much smaller sealing zone structures of +/+ and RQ/+ OCLs (Figure 5B). Similarly-large cells were observed also when cells were plated on plastic (Figure 6B below). Importantly, +/+, RQ/+, and RQ/RQ OCLs express similar levels of SNX10 protein (Figure 5 – figure supplement 1), indicating that the R51Q mutation does not affect overall SNX10 protein levels. cDNA sequencing revealed that +/+ OCLs express wild-type *Snx10* mRNA and RQ/RQ OCLs express mutant mRNA exclusively, while OCLs from RQ/+ mice express both *Snx10* mRNA species (Figure 5 – figure supplement 2).

**Figure 5:**
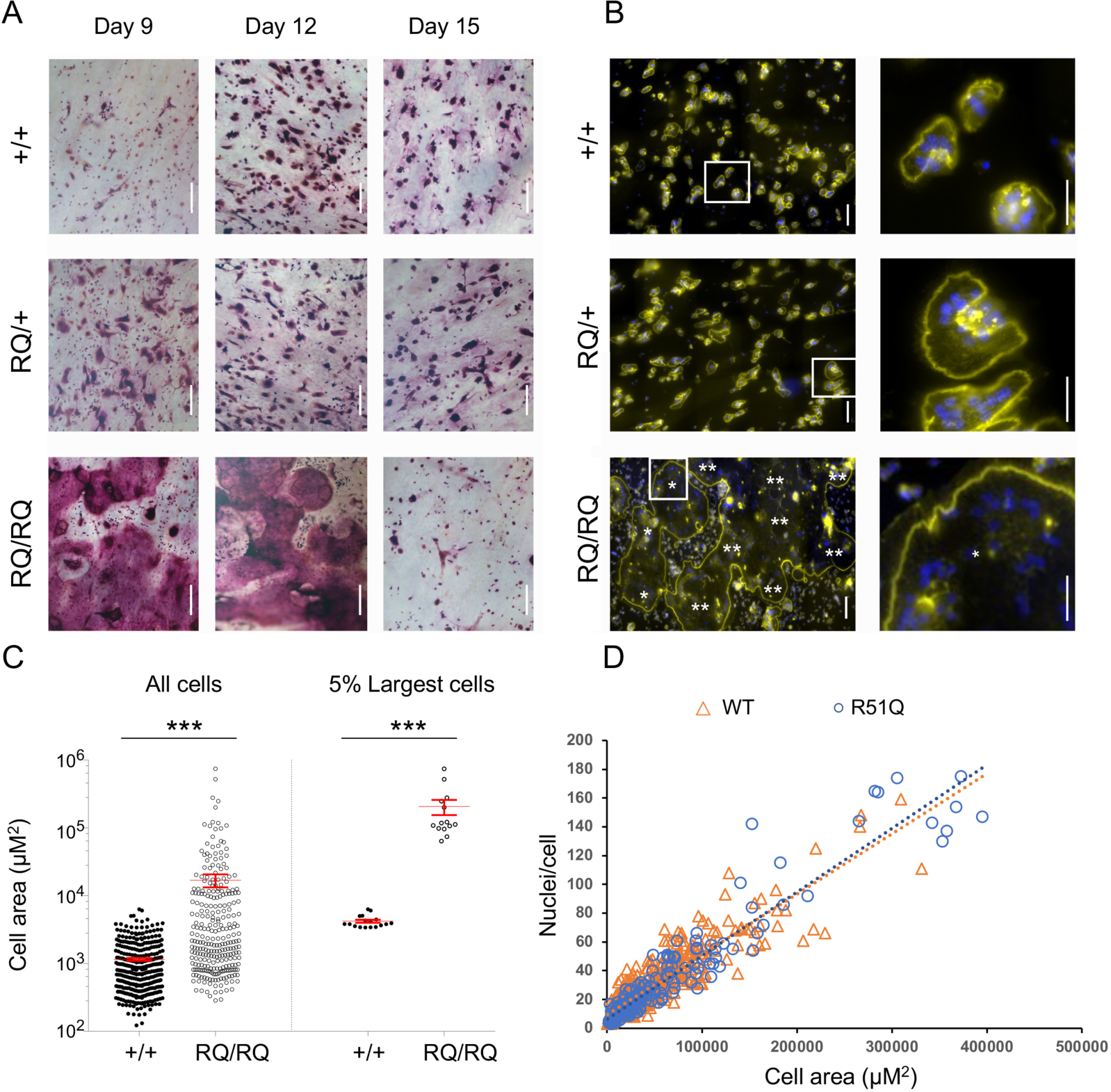
Giant osteoclasts from R51Q SNX10 homozygous mice. Spleen cells from +/+, RQ/+, and RQ/RQ mice were seeded on slices of bovine bone and grown with M-CSF and RANKL for up to 15 days. **(A)** Bone fragments were stained for TRAP. Scale bars: 200 μm. Figures are representative of a total of 4 +/+, 2 RQ/+, and 9 RQ/RQ mice analyzed in three experiments. Image of RQ/RQ cells at day 12 is a composite of two identical pictures taken at different focal planes due to unevenness of the bone surface. **(B)** Cells were stained with phalloidin (yellow; actin in sealing zones) and Hoechst 33258 (blue; DNA in nuclei). Left: Scale bar: 100 μm. Most of the RQ/RQ image is occupied by two giant cells, marked with one or two asterisks, respectively. Each figure is a composite of 15 smaller contiguous fields. Right: Magnified views of boxed areas in the left panels; asterisk in RQ/RQ image indicates a region within a single OCL. Scale bar: 50 μm. Figures are representative of a total of 3 +/+, 3 RQ/+, and 3 RQ/RQ mice analyzed in two experiments. **(C)** RQ/RQ OCLs are dramatically larger than +/+ OCLs. Left: The average size (mean±SD) of OCLs was 1,151±54 μm^2^ (+/+; N=353 cells) vs. 16,892±3,682 μm^2^ (RQ/RQ; N=280 cells). Right: The average sizes of the largest 5% of OCLs in either genotype was 4,207±217 μm^2^ (+/+; N=17 cells) vs. 206,175±52,323 μm^2^ (RQ/RQ; N=14 cells). ***: P<0.0001 by Student’s t-test. Note that the Y axis scale is logarithmic. Data were collected from a total of 2 +/+ and 5 RQ/RQ mice from two experiments. **(D)** RQ/RQ and +/+ OCLs exhibit similar nuclear densities. Scatter diagram depicting the number of nuclei and area per cell in RQ/RQ OCLs (N=205 cells) and +/+ (N=544 cells) OCLs of similar size ranges. Linear trendlines for both genotypes overlap (R^2^: +/+ =0.8143, RQ/RQ=0.9095). Cells were grown on plastic plates. See also Figure 5 – figure supplements 1, 2, and 3.

**Figure 6:**
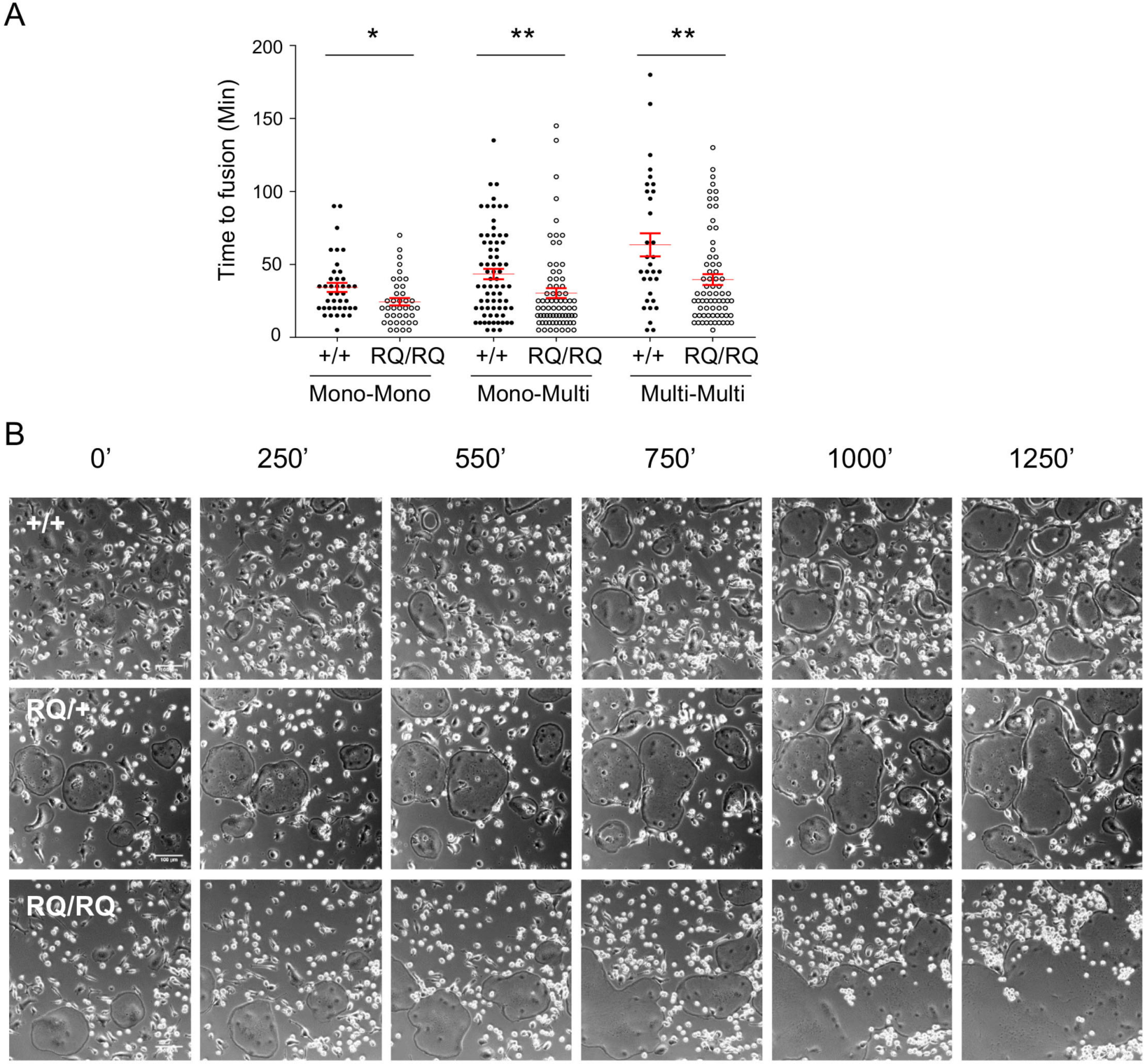
Fusion dynamics of homozygous R51Q SNX10 OCLs. (**A**) Time elapsed between initial formation of contact and subsequent fusion of cells. Fusion categories are between two mononuclear monocytes (Mono-Mono), between a mononuclear monocyte and a multinuclear OCL (Mono-Multi) and between two multinuclear OCLs (Multi-Multi). Data is presented as mean±SE, N=31-73 fusion events per category and genotype. *: P=0.017, **: P<0.0085 by Student’s t-test. Data were obtained from 3 +/+, 3 RQ/+, and 7 RQ/RQ mice in three experiments. (**B**) Images captured from phase live cell imaging videos showing fusion of +/+, RQ/+, and RQ/RQ OCLs at the indicated times (Videos 1, 2, and 3 for +/+, RQ/+, and RQ/RQ cells, respectively). Cells were grown on plastic tissue culture plates. Scale bar = 100 μm. Composite videos of 3×3 fields showing larger field views of fusion of +/+ and RQ/RQ cells are presented as Videos 4 and 5, respectively. Small bright cells are monocytes that have not yet fused.

Measurements of the areas of individual OCLs grown on bone fragments indicated that the average size of RQ/RQ OCLs was 14.6 times larger than +/+ OCLs; this ratio increased to 49 when the largest 5% of cells of each genotype were compared (Figure 5C). No similarly-large cells were observed in +/+ or RQ/+ cultures. Large OCLs could arise from increased spreading of the cells and/or from increased fusion of precursor cells. In order to examine this issue, we plotted the number of nuclei in individual +/+ and RQ/RQ OCLs as a function of their size (Figure 5D). Both parameters varied considerably among individual OCLs within each genotype. However, the relationship between nuclear number and cell size was modeled very well by a linear approximation in OCLs from both genotypes (Figure 5D), most likely due to the flatness of OCLs. The linear approximations for both genotypes shown in Figure 5D overlapped, indicating that in both genotypes the contribution of adding individual cells to OCL area was the same. The correlation between nuclear number and cell area was maintained also in the exceptionally-large RQ/RQ OCLs (Figure 5 – figure supplement 3). We conclude that giant RQ/RQ OCLs are most likely produced through increased fusion of precursor cells.

### Giant RQ/RQ OCLs are produced by continuous fusion of precursor cells

In order to better understand the basis for the abnormal fusion process that occurs in R51Q SNX10 OCLs, we examined the dynamics of the fusion process by live cell imaging. In culture, production of OCLs initiates when two mononuclear monocytes fuse, and the resulting binucleated cell continues to grow mainly by successive fusion with additional mononuclear cells (Levaot et al., 2015; Soe et al., 2015). In these cases, the cells come into close proximity with each other, juxtapose their membranes, and fuse after 30-60 minutes (+/+ cells, Figure 6A). Multinucleated cells tend not to fuse with each other, and the majority of interactions between multinucleated cells end in separation (Levaot et al., 2015; Soe et al., 2015). Examination of this process in RQ/RQ cells revealed that the time that elapsed between establishing of physical contact between the cells and their subsequent fusion was significantly shorter than in +/+ cells (Figure 6A). More rapid fusion of RQ/RQ cells was observed in all three fusion modalities – fusion between two mononuclear cells, between a mononuclear cell and a multinuclear cell, and between two multinuclear cells. In agreement with previous studies (Levaot et al., 2015; Soe et al., 2015), fusion events between two multinuclear cells were rare in +/+ cultures. Most interactions between two adjacent multinuclear +/+ OCLs ended in separation after a relatively long time (393.8±56.2 minutes, Mean±SE, N=59 events). In stark contrast, *all* cases in which multinucleated RQ/RQ cells were found adjacent to each other led to rapid fusion in the times noted in Figure 6A. No cases were observed in which adjacent multinuclear RQ/RQ cells separated without fusion, indicating that fusion, not separation, is the predominant outcome of interactions between multinuclear RQ/RQ cells.

The OCL fusion process typically ends when a multinuclear OCL becomes surrounded by other multinuclear cells with which it does not fuse, or when mononuclear fusion partners are no longer available. Fusion of +/+ and of RQ/+ OCLs proceeded in this manner and terminated when the entire field of view was occupied by individual OCLs arranged side-by-side in a dense pattern (Figure 6B,+/+ and RQ/+ cells, t=1250 minutes; Videos 1, 2, 4). In contrast, multinuclear RQ/RQ OCLs continued to fuse with each other, generating ever-larger OCLs that continued to grow by fusing with other multinuclear OCLs. In these cases, the fusion process ended when all available cells had fused into a single large cell that was often larger than the entire field of view, or when the cell died (Figure 6B, RQ/RQ cells; Videos 3, 5). We conclude that the R51Q SNX10 OCL phenotype is caused by specific abnormalities in key features of the fusion process relative to +/+ cells: RQ/RQ cells fuse more rapidly, fusion between multinuclear cells is extremely common, and the fusion process continues unabated to form extremely large OCLs.

### RQ/RQ osteoclasts exhibit abnormal ruffled borders and are severely functionally impaired

We next examined the structure of RQ/RQ OCLs grown on bone *in vitro* by transmission electron microcopy (TEM). Ruffled borders, the basal membranal structures through which the cells secrete acid and proteases onto the bone surface below, were present in all +/+ and RQ/+ cells, visible as areas in which heavily convoluted membranous folds extended from the ventral aspect of the cells onto and into the bone surface below (Figure 7A). In contrast, RQ/RQ OCLs lacked ruffled border structures. Some mutant cells did not appear to make contact with bone, while others made contact via a nearly flat surface that completely lacked the convoluted structure of the ruffled border membranes (Figure 7A). Similar findings were made when the structures of endogenous OCLs in femurs of +/+ and RQ/RQ mice were examined (Figure 7B).

**Figure 7:**
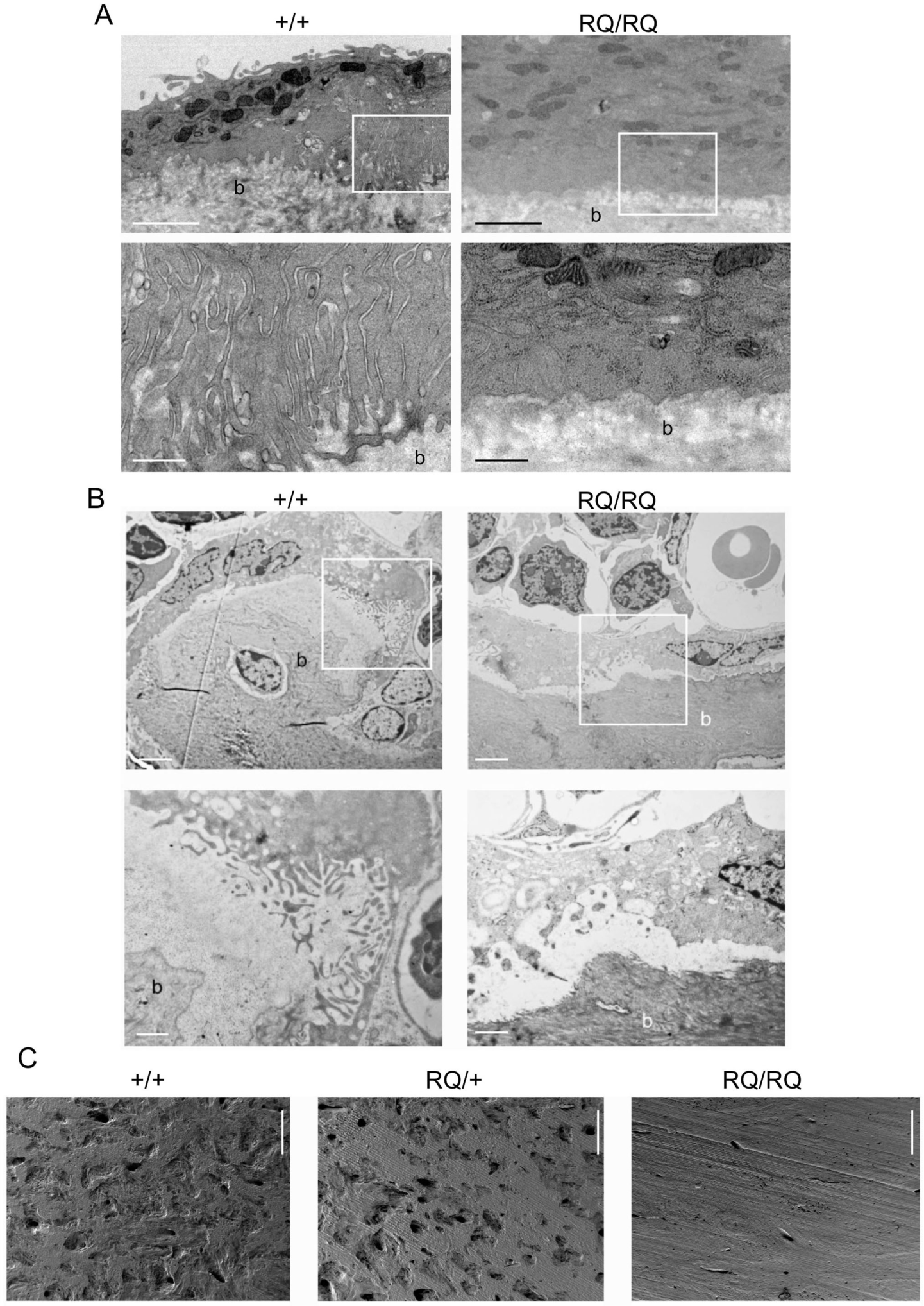
Giant OCLs of R51Q SNX10 homozygous mice exhibit structurally-abnormal or missing ruffled borders. Spleen cells from +/+ and RQ/RQ mice were seeded on slices of bovine bone and grown in the presence of M-CSF and RANKL for 15 days. **(A)** Transmission electron microscopy (TEM) images of +/+ and RQ/RQ OCLs grown on bone. Top: Scale bars: 2 μm. Bottom: Enlarged images of boxed regions in top panels. Scale bar: 500 nm. RQ/+ cells were indistinguishable from +/+ cells. Images are representative of a total of 17 +/+, 5 RQ/+, and 17 RQ/RQ cells examined from one mouse per genotype. b, bone **(B)** Similar to A, showing sections of mouse bones with endogenous OCLs. Top: Scale bars: 10 μm. Bottom: enlarged images of boxed regions in top panels. Scale bars: 1 μm. b, bone. Images shown are representative of 6-10 images obtained from two female mice aged 5 weeks, per genotype. **(C)** RQ/RQ OCLs are inactive. Scanning electron microscopy (SEM) images of bovine bone fragments seeded with +/+, RQ/+, or RQ/RQ cells as indicated. Scale bar: 200 μm. Images are representative of data collected from 5 +/+, 3 RQ/+, and 9 RQ/RQ mice in three experiments.

The large size and absence of ruffled border structures of RQ/RQ OCLs prompted us to examine their capacity to resorb bone. To this end we seeded +/+, RQ/+, and RQ/RQ splenocytes on fragments of bovine bone and grew them for 15 days in the presence of M-CSF and RANKL. Cells from all three genotypes grew well and fused as indicated in Figure 5A. We then removed the cells and examined the formation of resorption pits on the bone surface, which indicate OCL activity, by scanning electron microscopy (SEM). As seen in Figure 7C, OCLs from +/+ and RQ/+ mice readily formed resorption pits that were similar in size and in appearance. In contrast, RQ/RQ OCLs did not form pits at all, indicating they are severely deficient in function. Since the pit resorption assay provides an accumulated record of resorption activity throughout the life of the culture, these results indicate that the RQ/RQ cells were at no time capable of significant activity. We conclude that the massive osteopetrosis of homozygous R51Q SNX10 mice arises from virtual lack of OCL-mediated bone resorption, which is caused by a combination of the absence of ruffled border structures of RQ/RQ OCLs and by their large size and reduced lifespan. The latter effects are caused by severe abnormalities in the cell fusion processes in which these cells are produced.

## Discussion

Autosomal recessive osteopetrosis is a lethal genetic disease whose main symptoms are caused by OCL dysfunction. The only treatment currently available for ARO is hematopoietic stem cell transplantation, a highly-invasive procedure that is not suitable for all patients and which does not completely reverse the damage caused by the initial failure in bone resorption (Sobacchi et al., 2013; Tolar et al., 2004; Wu et al., 2017). These facts, together with the rare occurrence of ARO and its heterogeneous genetic basis, make development of accurate mouse model systems for ARO essential for clarifying its cellular and molecular bases, and for developing alternative approaches for diagnosis and therapy. In this study, the mechanism underlying the development of ARO caused by the R51Q mutation in the SNX10 protein (Aker et al., 2012) is explored.

### The SNX10 R51Q mutation leads to a unique form of ARO that combines reduced OCL numbers and impaired activity

Homozygous R51Q SNX10 mice exhibit multiple phenotypes that closely mimic the clinical manifestations in humans bearing the corresponding mutation. A key feature among these is massive, early-onset, and widespread osteopetrosis that is caused by dysfunctional OCLs. Additional similarities include missing and impacted teeth, occasional osteomyelitis, stunted growth, failure to thrive, and a significantly reduced lifespan (Sobacchi et al., 2013; Tolar et al., 2004; Wu et al., 2017). Hepatosplenomegaly and anemia, which often develop in ARO patients, were not found in RQ/RQ mice, possibly due to their short lifespan that precludes development of these symptoms. These findings confirm the R51Q SNX10 mutation as a causative factor in ARO.

A prominent feature of RQ/RQ OCLs is their size. These cells fuse rapidly and continuously, growing to become dramatically larger than wild-type OCLs, which are large cells in their own right. The size phenotype is likely cell-autonomous since it is detected after several days of differentiation *ex vivo* in culture, and occurs irrespective of the tissue origin of the monocyte precursor cells (spleen, bone marrow, or circulating monocytes) or of the surface upon which they are grown. Another striking feature of RQ/RQ OCLs is their inability to resorb bone *ex vivo*, which is most likely in part the result of structurally-defective or absent ruffled borders. All the previously-known proteins whose mutations cause ARO by inhibiting OCL activity localize to secretory vesicles in OCLs, and many of these mutations have been shown to impair ruffled border formation (Sobacchi et al., 2013). SNX10 localizes to endosomes; its R51Q mutation that is located in the membrane-binding PX domain might prevent proper localization of SNX10 and lead to aberrant ruffled border structure. Alternatively, SNX10 associates with the ATP-dependent vacuolar proton pump of OCLs (V-ATPase) and affects its function (Chen et al., 2012), suggesting the R51Q mutation might induce ARO by mis-localizing the proton pump and affecting acidification. An intriguing possibility is that the size of RQ/RQ OCLs, which increases their fragility *in vitro* and reduces their abundance *in vivo*, renders the functional lifespan of these cells too short to perform significant bone resorption. Further studies are required to clarify these issues.

Two major types of ARO have been described in humans and in mouse models for this disease: OCL-poor ARO, which is caused by mutations that block production of OCLs, and OCL-rich disease, in which normal or increased numbers of inactive OCLs are present due to mutations that reduce OCL activity (Sobacchi et al., 2013). The R51Q SNX10 mutation defines a third category, in which OCLs are both non-functional and their abundance is significantly reduced *in vivo*. We believe that the bases for this unique phenotypic combination are the dual consequences of the R51Q mutation in SNX10, which exerts significant effects on osteoclast maturation and survival, and renders the cells inactive.

### Differential effects of SNX10 mutations in humans and in murine models

Based on the assumption that the R51Q mutation in SNX10 is a “loss of function” mutation, SNX10-KD mice were initially used as a model for ARO (Ye et al., 2015). While osteopetrosis and other features of the RQ/RQ phenotype that are described here were also detected in SNX10-KD mice, significant differences exist between these models. The SNX10-KD mouse phenotype is characterized by an increased mass of bone that exhibits, paradoxically, markedly reduced mineral content (Ye et al., 2015). This rachitic phenotype is attributable to disruption of SNX10 function in the stomach that leads to higher gastric pH, abnormal stomach appearance, and reduced absorption of dietary calcium (Ye et al., 2015). The femoral mineral content of RQ/RQ mice is reduced, but to a lesser degree than that seen in SNX10-KD mice. Moreover, the main defects that give rise to the rachitic phenotype of SNX10-KD mice do not exist in RQ/RQ mice: stomach appearance and gastric pH are normal, as are serum levels of PTH and, critically, calcium. We note that concentrations of circulating calcium are normal in humans suffering from ARO induced by R51Q SNX10 (Figure 3F) and by the Vasterbotten frameshift mutation in SNX10 [10]; rickets symptoms were not reported in these cases (Aker et al., 2012; Stattin et al., 2017). Administering calcium gluconate, which bypassed reduced calcium resorption in SNX10-KD mice and resolved their rachitic phenotype (Ye et al., 2015), is therefore not a general treatment for all forms of SNX10-based ARO.

Differences in OCL morphology were also noted between the RQ/RQ mice and other ARO patients and mouse models. Specifically, OCLs two to five times larger than controls were reported in cultures prepared from blood monocytes of patients of the genetically-distinct SNX10 Vasterbotten variant of ARO (Stattin et al., 2017), and in mice lacking OSTM-1 (Pata and Vacher, 2018) that model another cause of ARO. The size of these OCLs is significantly smaller than RQ/RQ OCLs, indicating that the cellular consequences of those mutations differ from R51Q SNX10. Large OCLs were not described in mice carrying whole-body or OCL-specific knockdowns of SNX10 (Ye et al., 2015). Large OCLs were also not noted in reports of SNX10 knockout mice, although the genetic manipulation of the *Snx10* gene in these mice has not been described (Zhou et al., 2016). Moreover, heterozygosity for R51Q SNX10 does not elicit OCL or bone phenotypes, making it unlikely that the R51Q SNX10 mutant protein acts through dominant gain-of-function or dominant-negative mechanisms. We thus propose that R51Q SNX10 acts by a loss-of-function mechanism, or by a recessive gain-of-function mechanism that becomes evident only in the absence of the wild-type protein.

### OCL size is limited by an active cellular mechanism

The dynamics whereby precursor cells of the monocyte lineage initiate the fusion process and then continue to fuse, forming multinucleated OCLs, have been described in great detail (Levaot et al., 2015; Soe et al., 2015). However, the regulation of this process and, in particular, how fusion is arrested when cells reach “maturity”, are mostly unknown. The fusion process in cultures of wild-type monocytes, stimulated by RANKL, halts when multinucleated OCLs either run out of fusion partners or when they become surrounded by other large multinucleated OCLs, with which they tend not to fuse ((Levaot et al., 2015; Soe et al., 2015); e.g., Video 4). In wild-type cultures, this transition from the fusing to the arrested state leads to accumulation of “mature” OCLs with predictable ranges of sizes and numbers of nuclei that rarely exceed 200,000 µm^2^ and 100 nuclei, respectively, per cell (Figure 5D). In contrast, RQ/RQ OCLs continue to fuse with each other, forming giant heterokaryons that can become orders of magnitude larger than a mature wild-type cell and may contain several hundreds of nuclei. The present study, in which a defined mutation in a specific gene product leads to continuous fusion and abolishes control of OCL size, clearly indicates that OCL size is actively regulated by cellular-genetic mechanisms. The data also provide compelling evidence that SNX10 participates in this mechanism by promoting the fusion-terminating process. The R51Q mutation disrupts this role of SNX10 and enables fusion to continue unabated, leading to the development of giant, unstable, and non-functional OCLs.

Reduced production of OCLs negatively affects their activity, a fact that has been utilized clinically in the form of drugs, such as the anti-RANKL antibody Denosumab, that inhibit bone resorption by down-regulating OCL production. The present study indicates that excessive fusion that produces aberrantly-large OCLs also negatively affects OCL survival and function. Optimal OCL activity is therefore associated with a defined size range, suggesting that promotion of excessive fusion that makes these cells too large and fragile might be a distinct strategy for reducing OCL-mediated bone resorption in disease.

Multiple mutations in SNX10 have now been described in ARO (Aker et al., 2012; Pangrazio et al., 2013; Stattin et al., 2017). Our study underscores the importance of mimicking these human mutations as closely as possible in preclinical models, since loss-of-function approaches do not necessarily generate the complex phenotypes seen at the cellular level and prevent examination of possible gain-of-function effects of these mutations.

### Materials and Methods

#### Animal studies

##### Generation of R51Q SNX10 knock-in mice

Mice carrying the R51Q mutation in SNX10 were constructed by CRISPR at The Weizmann Institute. Candidate sgRNA sequences were selected from SNX10 genomic sequences using sgRNAScorer 2.0 (https://crispr.med.harvard.edu/sgRNAScorer/;(Chari et al., 2015)) and the CRISPR design tool of the Zhang lab, MIT (http://crispr.mit.edu/). The sequence 5’-ACACCAGCAACGCGTTGCTT-3’ (non-coding strand) was used. Linearized templates were transcribed with the MEGAshortscript T7 kit (sgRNA) or the mMessage Machine T7 kit (Cas9 mRNA), and purified by the MEGAclear purification kit according to the manufacturer’s instructions (all from Ambion-ThermoFisher Scientific, Waltham, MA). The following single-stranded DNA oligomer (non-coding strand; mutated sequence) was synthesized by Integrated DNA Technologies (Skokie, IL) used for homologous recombination without purification: *GAACTAGACTCATGGCTCCCGGTAATTTAAACTTTAAGTCACTT*<u>*AC*ACCAGCAACGCGTTaCTc</u>TGGAGTCTTTGCCTCAGCCACACGAAtTCTCTATACCTTCTTtGTACACAAGATGTTTTCATTG TAAAACACATGCTATTTGT*CTGTATAGAAACAGAAAAAA* Non-italicized nucleotides are from coding sequences of exon 4 of the SNX10 gene. Shown are the guide sequence (underlined), and four mutations introduced by this oligo into genomic DNA (lowercase, underlined, in 5’ to 3’ order): G>A, T>C (silent mutations to prevent re-cleavage); C>T (silent, introduces an EcoRI site for genotyping); C>T (R51Q mutation).

Cas9-encoding RNA (100 ng/ul), sgRNA (50 ng/ul), and the DNA oligomer (200 ng/ul), dissolved in water, were co-injected into the cytoplasm of fertilized oocytes obtained from C57BL/6JOlaHsd mice, and the injected oocytes implanted in pseudo-pregnant foster mothers. Potential founder mice were identified by PCR of tail biopsy DNA (see below), and two (founders 43 and 87) were bred into independent lines by crossing with B6(Cg)-Tyr^c-2J^/J (albino C57Bl/6J) mice. The two mutant alleles are designated by the Mouse Genomic Nomenclature Committee (MGNC) as Snx10^em1AE1^ (line 43) and Snx10^em2AE1^ (line 87), but are referred to here as RQ for convenience.

Potential off-target sites for the sgRNA were selected using the CRISPR design tools of the Zhang lab, MIT (http://crispr.mit.edu/ or https://benchling.com/crispr). The five off-target sites that scored highest were selected (sequences: acagaaccaacgccttgcttcggg (Chr. 13), agaacagtaacgcgctagcttggga (Chr. 2), agtaaagcaacgctttgctttggt (Chr. 12), ataccataaacacgttgctttag (Chr. 7) and tcaccaggaagtcgttgcttaag (Chr. 13). DNA segments of 450-500 bp in length, centered around these sequences, were amplified by PCR and sequenced; no mutations were detected. Founders were bred with wild-type B6(Cg)-Tyr^c-2J^/J mice, and heterozygous F1 offspring were separately bred with B6(Cg)-Tyr^c-2J^/J mice. Heterozygous F2 offspring of a single F1 mouse were intercrossed to generate wild-type (+/+), heterozygous (+/RQ), and homozygous (RQ/RQ) progeny.

##### Genotyping

Genotyping by PCR was performed using oligos int3FW (5’-ACATGCCCACTGGAGTTTTC-3’ and int4REV (5’-CCTGTGCCCTGTTCTTACCT-3’) in a program that included of 35 cycles of denaturation (95°C, 30 seconds), annealing (58°C, 30 seconds) and elongation (72°C, 30 seconds). Cleavage of the mutant PCR product with EcoRI yielded fragments of 387bp and 142bp, while the wild-type allele was not cut.

##### Animal Husbandry

At both the Weizmann Institute and at the University of Ulm, mice were housed in a barrier facility kept at 22±2°C on a light/dark cycle of 12h:12h, with food and water provided *ad libitum*. Colonies were maintained by intercrossing RQ/+ mice as described above to produce +/+, RQ/+, and RQ/RQ littermate mice. All pups were kept with their mothers until weaning (4 weeks), after which teeth-bearing mice (+/+, RQ/+) were separated by sex and housed up to 5 per cage. Toothless RQ/RQ mice were left to nurse up to 6-8 weeks of age, in the presence of younger litters to ensure continued lactation. All experiments comparing mice (e.g., body weights, lengths, CTX assays) were performed on sex-matched littermate mice prior to weaning. Primary OCLs were prepared from mice of either sex either just before or after weaning.

All experiments performed at the Weizmann Institute were approved by the Weizmann Institute IACUC, and were conducted in accordance with Israeli law. All experiments performed at the University of Ulm were approved under animal license #1329 of the state of Baden-Württemberg and in accordance with local law.

### Human Studies

Retrospective blood calcium levels from R51Q SNX10 patients were obtained from their files at the Hadassah Medical Center, Jerusalem. Researchers conducting this study were blinded to the identity of the patients. Access to patient data was approved by the Ethics Committee of the Hadassah Medical Center and by the Institutional Review Board of The Weizmann Institute.

### Histomorphometry, histology, and skeletal analyses

#### Calcein labeling

Mice were injected IP with 30 mg/kg body weight of calcein solution (10 mg/ml calcein in 0.15M NaCl/2% NaHCO_3_), on days 1 and 8, and were sacrificed on day 10. Following removal pf skin and internal organs, the mice were fixed in 4% paraformaldehyde/PBS for 3 days, and then kept in 70% ethanol until analyzed. Mice were analyzed prior to weaning at 20-25 days of age; both male and female mice from both R51Q SNX10 mouse lines were analyzed.

#### X-ray imaging

Mouse skeletons were scanned using an MX20 Cabinet X-Ray system (Faxitron X-Ray, Lincolnshire, IL) instrument using standard settings.

#### Micro-CT measurements

Mouse skeletons were scanned using a SkyScan1176 µCT (Bruker, Billerica, MA) instrument using a 50µm aluminum filter, 9µm voxel size and a 1° rotation step. The software programs NRecon, Data Viewer, CTAn, CTVol (Bruker) were used for analysis. For tissue mineral density measurements the bones were rehydrated in PBS and phantoms of 0.25g and 0.75g CaHA/cm^3^ (Bruker) were scanned the same day for calibration.

#### Bone histomorphometry

For bone formation rate measurements, osteoblast/osteocyte count, and von-Kossa staining, non-decalcified bones were embedded in methyl-methacrylate. Bone formation rate was measured on 8µm sections after embedding with DAKO fluorescence mounting medium (Agilent, Santa Clara, CA). For osteoblast/osteocyte analysis 5µm sections were deplasticized and stained with 0.1% toluidine blue in 0.1M sodium acetate buffer (pH 5.0). For osteocytes, the number of nuclei inside trabecular bone were counted. For osteoblasts, large square cells in contact with the bone and possessing soft edges were included in the count and their surface in contact with bone was marked. For TRAP staining bones were decalcified in 15% EDTA, pH 8.0, for 14 days and embedded in paraffin. 5µm sections were stained for TRAP using a leukocyte acid phosphatase kit (Sigma-Aldrich, St.Louis, MO). Slides were counterstained with hematoxylin and large, flat TRAP positive cells in contact with trabecular and cortical bone were counted. Analysis of all slides was carried using OsteomeasureXP (OsteoMetrics, Decatur, GA; version 1.01)

#### Histology

Mouse tissues were fixed in 10% formalin, embedded in paraffin, and sectioned at 4 microns. Samples containing bone were decalcified prior to embedding. Sections were stained with Hematoxylin Solution, Gill no. 3 and with Eosin Y solution, according to the manufacturer’s instructions (Sigma). Mice analyzed were matched for age and sex.

### Measurement of bone markers in blood

#### CTX

Was measured using the Ratlaps CTX-1 EIA kit (Immunodiagnostic Systems (IDS), Tyne & Wear, UK) according to the manufacturer’s instructions. Plasma samples were prepared from blood collected from the orbital sinus from 23-day old pups that had been fasted for 6 hours.

#### PTH

Was measured using a PTH EIA kit (Ray Biotech, Norcross, GA) according to the manufacturer’s instructions. Plasma samples were prepared from blood collected from the orbital sinus from non-fasted 25-day old pups.

#### 25-(OH)-vitamin D3

Was measured using the 25-(OH)-vitamin D3 kit Enzo (Farmingdale, NY) according to the manufacturer’s instructions. Samples were the same ones used for PTH determination.

#### P1NP

Was measured using the Rat/Mouse PINP EIA kit (IDS) according to the manufacturer’s instructions. Plasma samples were prepared from blood collected via cardiac puncture from unfasted 5-week old male and female mice.

CTX, PTH, and 25-(OH)-vitamin D3 results were analyzed by four parameter logistic regression using SigmaPlot software (Systat Software, San Jose, CA). P1NP results were analyzed by Prism 6 software (GraphPad, San Diego, CA). Sex distribution of mice is noted in the relevant figure legends.

### Measurement of gastric pH

WT and RQ/RQ mice, between 20-30 days old and still nursing, were of both sexes and lines. Mice were fasted for two hours, weighed, sacrificed by decapitation, and stomachs were dissected from the esophagus to the pylorus and weighed. Stomach contents were then flushed with 500 microliters of water (pH 7.8), diluted with an equal amount of water, and pH of the mixture was measured with a pH meter.

### Culture of primary mouse osteoclasts and osteoblasts

#### Culture of primary mouse osteoclasts on plastic

Osteoclast precursors were isolated from spleens of mice aged 6-8 weeks. Spleens were dissociated into un-supplemented α-Minimal Eagle’s Medium (α-MEM; Gibco-Thermo Fisher Scientific or Sigma) Following lysis of erythrocytes with red blood cell lysis buffer, cells were seeded at a density of 5×10^6^ cells/well (for +/+ or RQ/+) and 2.5×10^6^ cells/well (RQ/RQ) in 6-well plates, or 2×10^6^ cells/well (+/+ or RQ/+) and 1×10^6^ cells/well (RQ/RQ) in 24-well plates. Cells were cultured in complete OCL medium (α-MEM supplemented with 10% fetal calf serum (FCS), 2 mM glutamine, 50 units/ml penicillin, and 50 g/ml streptomycin, as well as 20 ng/ml M-CSF (Peprotech, Rehovot, Israel) and 20 ng/ml RANKL (R&D Systems, Minneapolis, MN). Cells were incubated at 37 °C in 5% CO_2_ for 5-7 days with daily changes of medium.

#### Culture of primary mouse osteoclasts on bone fragments

Bovine femurs were sliced into small fragments with a diamond-tipped saw and stored in 70% ethanol at 4°C until use. Prior to seeding, bone slices were washed in PBS and placed in 24-well plates (2-3 slices/well) and incubated in OCL medium (without cytokines) for 1hr at 37 °C in 5% CO_2_. Cells were prepared as above and then seeded at a density of 1×10^5^ cells/well in 24-well plates and cultured in complete OCL medium that included M-CSF and RANKL as above, with changes of medium every 48-72 hours.

### Analysis of cultured OCLs

#### Cell staining, immunofluorescence, and cell size measurements

Bone: OCLs grown on bone slices were stained for TRAP using the Leukocyte Acid Phosphatase kit (Sigma). Due to the faster differentiation of mutant cells, they were stained 10 days after seeding while WT were stained 14 days. For actin staining, cells were permeabilized and fixed by incubating the slices in 0.5% Triton X-100/3% PFA for 3 minutes, followed by fixation in 3% PFA for 20 minutes and washes in PBS. Cells were then exposed to TRITC-phalloidin (diluted 1:500, Sigma) for 1 hour at room temperature. DNA was visualized by incubating the slices with Hoechst 33258 (diluted 1:3000, Molecular Probes, Eugene, OR) for 3 minutes.

Plastic: For measurements of nuclear number and cell size, M-CSF treated cells were seeded in µ-Slide 4 well imaging chambers, with a coverslip polymer bottom (Ibidi, Martinsried, Germany) at a density of 1×10^5^ cells/well and cultured in complete OCL medium supplemented with 20 ng/ ml M-CSF and 20 ng/ml RANKL for 4 days; medium was changed every 24 hours. Cells were washed with PBS, fixed with 3% paraformaldehyde in PBS for 15 min and then permeabilized with 0.5% Triton X-100 in 3% paraformaldehyde for 3 minutes. Cells were then stained with phalloidin and Hoechst 33258 as above. For quantification, OCL boundaries were manually marked using Image J software. Image analysis was done semi-automatically using Fiji software (Schindelin et al., 2012).

#### Live cell imaging

For time-lapse videos, cells were first cultured in complete OCL medium supplemented only with 20 ng/ml M-CSF. When confluent, cells were detached with Trypsin-EDTA and seeded at a density of 1×10^5^ cells/well in 24-well plates or on glass coverslips. Cells were then cultured in complete OCL medium supplemented with 20 ng/ml M-CSF and 20 ng/ml RANKL, with daily changes of medium for 3-4 days prior to imaging. Time-lapse images were acquired with an automated inverted microscope (DeltaVision Elite system IX71 with Resolve3D software modulus; Applied Precision, Inc., GE Healthcare, Issaquah, WA) using a 10×/0.30 air objective (Olympus, Tokyo, Japan). The microscope is equipped with an environmental box kept at 37 °C with a 5% CO_2_ humidified atmosphere. Images were acquired every 5 min, for up to 24 h. For image montages, 3×3 adjacent fields with 10% overlap were selected.

#### Live cell Imaging analysis

Image display and analysis were performed using Fiji (Schindelin et al., 2012). For montage display we used the stitching plugin (Preibisch et al., 2009). Some images were corrected for background and shading using the BaSic tool (Peng et al., 2017). Time (=number of video frames taken at 5 minute intervals) from first contact of cells to their ultimate fusion was determined from live cell imaging videos. Cells were followed backwards in time from fusion (defined as the time point where the cells shared a continuous stretch of cytoplasm the size of a diameter of a nucleus) to the time when they first made contact. Time between first contact and separation without fusion was measured in the same manner starting from cells that had just separated. Number of visible nuclei in each cell was annotated.

### Culture of primary mouse osteoblasts from calvarias

Osteoblast precursors were isolated from calvarias of 5-day old pups (both sexes combined). Dissected calvariae were placed in ice cold PBS containing 100 units/ml penicillin and 100 μg/ml streptomycin, and digested five times for 10 min each at 37°C in 1ml sterile digestion solution (α-MEM medium with penicillin/streptomycin as above, 1% Collagenase A (Roche, Basel, Switzerland) and 0.1 % Dispase II (Roche)). The first digestion fraction was discarded and fractions 2-5 were pooled in 500µl FCS on ice. Cells were seeded 6-well plates and grown in culture medium (α-MEM, 10% FCS, penicillin/streptomycin as above). Cells from each well were split into a 10cm dish 1 or 2 days later. For osteogenic induction, cells were seeded at 50000 cells/ml and induced at no more than 80% confluency the next day in culture medium supplemented with 100 µg/ml ascorbic acid and 1 mg/ml β-glycerophosphate, or left untreated as control. The medium was changed every 3-4 days.

### Analysis of cultured osteoblasts

Early mineralization was assessed quantitatively by alkaline phosphatase (ALP) staining (Amplite colorimetric alkaline phosphatase kit; AAT Bioquest, Sunnyvale, CA) after 7 days of culture. ALP staining was assessed qualitatively by fixation of cells with 4% PFA and staining with Fast Violet B salt. Late mineralization was followed by fixation with 4% PFA cells and staining with 1% Alizarin Red (pH 4.2). Quantification was carried out by measuring absorbance at 570nm after dissolving the dye with 100mM cetylpyridinium chloride.

### Electron microscopy analyses

#### Scanning Electron microcopy (SEM), pit resorption assay

After culturing osteoclasts on bone slices for 15-17 days as described above, cells were removed with cotton swabs and 1% bleach. The bone fragments were washed, dehydrated through a graded ethanol solution series, and dried. The samples were then mounted on aluminum stubs, sputter-coated with iridium (CCU-010, Safematic, Bad Ragaz, Switzerland), and visualized with a Zeiss Sigma 500 scanning electron microscope (Zeiss, Oberkochen, Germany).

#### Transmission electron microscopy (TEM)

Bone slices carrying OCLs that had been grown for 10-11 days were fixed overnight in 3% paraformaldehyde, 2% glutaraldehyde in 0.1 M cacodylate buffer containing 5 mM CaCl_2_ (pH 7.4) at 4°C. The bones were decalcified in an EDTA solution (1.9% glutaraldehyde, 0.15M EDTA in 0.06M sodium cacodylate buffer) for 3 weeks, postfixed in 1% OsO4 in water followed by post-post fixation in 2% uranyl acetate. The specimens were then dehydrated in graded ethanol solutions and embedded in Agar 100 epoxy resin (Agar Scientific Ltd., Stansted, UK). Ultrathin sections (70-90 nm) were viewed and photographed with an FEI Tecnai SPIRIT transmission electron microscope (FEI, Eindhoven, The Netherlands), operated at 120 kV and equipped with an EAGLE CCD camera.

### Protein blot analysis

Cells were washed once in PBS and lysed in RIPA buffer (50 mM Tris (pH 8), 150 mM NaCl, 1% Nonidet P-40, 0.5% Deoxycholate, 0.1% SDS), supplemented with protease inhibitor cocktail (1 mM N-(α-aminoethyl) benzene-sulfonyl fluoride, 40 uM bestatin, 15 uM E64, 20 uM leupeptin, 15 uM pepstatin; Sigma). Protein concentrations were determined by the Bradford method (Bio-Rad protein dye reagent; Biorad, Hercules, CA) using bovine serum albumin as a standard. SDS-PAGE and protein blotting were performed as described (Gil-Henn and Elson, 2003). Primary antibodies used included antibodies to actin (Sigma, clone AC-40) and SNX10 (Sigma, and Abcam, Cambridge, UK). Enhanced chemiluminescence signals were visualized using an Imagequant LAS 400 Mini instrument (GE Healhtcare Biosciences, Uppsala, Sweden) and quantified (GelPro Analyzer V.4, Media Cybernetics,Rockville, MD)

### Quantification and Statistical Analysis

Data were analyzed by ANOVA, two-tailed Student’s t-test or the Mann-Whitney U-test as indicated. Number of samples and their definition as well as data errors are indicated in each figure. In studies where individual mice or groups of mice were compared, each mouse represented a biological repeat. In studies where behavior of individual cells was compared, each cell represented as a biological repeat and the number of mice from which the cells were obtained is indicated in the figure legend. Analyses are based on all data collected except in Table 1, where PTH and Vitamin D data exclude outliers beyond two standard deviations from the sample mean. In most cases the phenotype of RQ/RQ samples was too strong to make blinding of sample identity practical. Statistics analyses were performed using Prism 6 (GraphPad) or JMP13 (SAS, Cary, NC). The level of statistical significance was set at P=0.05.

## Acknowledgements

We thank Dr. Timothy Dahlem and the University of Utah Mutation Generation and Detection Core Facility for reagents. We also thank Dr. Rebecca Haffner-Kraus, Ms. Golda Damari, Dr. Alina Berkovitz, and Ms. Sima Peretz of the Weizmann Institute’s Transgenic and Knockout Core Facility for help in preparing the mouse models used in this study; Ms. Ofira Higfa and Mr. Neriah Sharabi (Weizmann Institute), and Mr. Thomas Neidlinger and Ms. Birgit Widmann (Ulm University) for expert animal care; Dr. Elena Kartvelishvily, Dr. Smadar Zaidman, and Ms. Ilana Sabanay of the Weizmann Institute’s Chemical Research Support Department for assistance with electron microscopy studies; and Dr. Ofra Golani of the Weizmann Institute’s Life Sciences Core Facilities department for help with image analysis. We also thank Prof. Memet Aker for discussions at the start of this project. AE is the incumbent of the Marshall and Renette Ezralow Professorial Chair. BG is the Erwin Neter Professor in Cell and Tumor Biology.

## Video file legends

**Video 1:** Phase light microscopy live cell imaging of +/+ monocytes as they fuse into OCLs. Individual images were obtained at 5-minute intervals for a period of 21 hours.

**Video 2:** Phase light microscopy live cell imaging of RQ/+ monocytes as they fuse into OCLs. Individual images were obtained at 5-minute intervals for a period of 21 hours.

**Video 3:** Phase light microscopy live cell imaging of RQ/RQ monocytes as they fuse into OCLs. Individual images were obtained at 5-minute intervals for a period of 21 hours.

**Video 4:** Phase light microscopy live cell imaging of +/+ monocytes as they fuse into OCLs. Image is a montage of 3×3 fields and covers an area of 2 × 2 millimeters. Individual images were obtained at 5-minute intervals for a period of 21 hours.

**Video 5:** Phase light microscopy live cell imaging of RQ/RQ monocytes as they fuse into OCLs. Image is a montage of 3×3 fields and covers an area of 2 × 2 millimeters. Individual images were obtained at 5-minute intervals for a period of 21 hours.

## References

Aker, M., Rouvinski, A., Hashavia, S., Ta-Shma, A., Shaag, A., Zenvirt, S., Israel, S., Weintraub, M., Taraboulos, A., Bar-Shavit, Z., and Elpeleg, O. (2012). An SNX10 mutation causes malignant osteopetrosis of infancy. J Med Gen 49, 221–226.

Bruzzaniti, A., and Baron, R. (2006). Molecular regulation of osteoclast activity. Rev Endocr Metab Disord 7, 123–139.

Chari, R., Mali, P., Moosburner, M., and Church, G.M. (2015). Unraveling CRISPR-Cas9 genome engineering parameters via a library-on-library approach. Nat Methods 12, 823–826.

Chen, Y., Wu, B., Xu, L., Li, H., Xia, J., Yin, W., Li, Z., Shi, D., Li, S., Lin, S., Shu, X., and Pei, D. (2012). A SNX10/V-ATPase pathway regulates ciliogenesis in vitro and in vivo. Cell Res 22, 333–345.

Cullen, P.J. (2008). Endosomal sorting and signalling: an emerging role for sorting nexins. Nat Rev Mol Cell Biol 9, 574–582.

Del Fattore, A., Cappariello, A., and Teti, A. (2008). Genetics, pathogenesis and complications of osteopetrosis. Bone 42, 19–29.

Faden, M.A., Krakow, D., Ezgu, F., Rimoin, D.L., and Lachman, R.S. (2009). The Erlenmeyer flask bone deformity in the skeletal dysplasias. Am J Med Genet A 149a, 1334–1345.

Gil-Henn, H., and Elson, A. (2003). Tyrosine phosphatase-epsilon activates Src and supports the transformed phenotype of Neu-induced mammary tumor cells. J Biol Chem 278, 15579–15586.

Levaot, N., Ottolenghi, A., Mann, M., Guterman-Ram, G., Kam, Z., and Geiger, B. (2015). Osteoclast fusion is initiated by a small subset of RANKL-stimulated monocyte progenitors, which can fuse to RANKL-unstimulated progenitors. Bone 79, 21–28.

Neutzsky-Wulff, A.V., Karsdal, M.A., and Henriksen, K. (2008). Characterization of the bone phenotype in ClC-7-deficient mice. Calcif Tissue Int 83, 425–437.

Novack, D.V., and Teitelbaum, S.L. (2008). The osteoclast: friend or foe? Annu Rev Pathol 3, 457– 484.

Palagano, E., Menale, C., Sobacchi, C., and Villa, A. (2018). Genetics of Osteopetrosis. Curr Osteoporos Rep 16, 13–25.

Pangrazio, A., Fasth, A., Sbardellati, A., Orchard, P.J., Kasow, K.A., Raza, J., Albayrak, C., Albayrak, D., Vanakker, O.M., De Moerloose, B., Vellodi, A., Notarangelo, L.D., Schlack, C., Strauss, G., Kuhl, J.S., Caldana, E., Lo Iacono, N., Susani, L., Kornak, U., Schulz, A., Vezzoni, P., Villa, A., and Sobacchi, C. (2013). SNX10 mutations define a subgroup of human autosomal recessive osteopetrosis with variable clinical severity. J Bone Miner Res 28, 1041–1049.

Pata, M., and Vacher, J. (2018). Ostm1 bifunctional roles in osteoclast maturation: insights from a mouse model mimicking a human OSTM1 mutation. J Bone Miner Res. 33, 888–898.

Peng, T., Thorn, K., Schroeder, T., Wang, L., Theis, F.J., Marr, C., and Navab, N. (2017). A BaSiC tool for background and shading correction of optical microscopy images. Nat Commun 8, 14836.

Preibisch, S., Saalfeld, S., and Tomancak, P. (2009). Globally optimal stitching of tiled 3D microscopic image acquisitions. Bioinformatics 25, 1463–1465.

Qin, B., He, M., Chen, X., and Pei, D. (2006). Sorting nexin 10 induces giant vacuoles in mammalian cells. J Biol Chem 281, 36891–36896.

Schindelin, J., Arganda-Carreras, I., Frise, E., Kaynig, V., Longair, M., Pietzsch, T., Preibisch, S., Rueden, C., Saalfeld, S., Schmid, B., Tinevez, J.Y., White, D.J., Hartenstein, V., Eliceiri, K., Tomancak, P., and Cardona, A. (2012). Fiji: an open-source platform for biological-image analysis. Nat Methods 9, 676–682.

Sobacchi, C., Schulz, A., Coxon, F.P., Villa, A., and Helfrich, M.H. (2013). Osteopetrosis: genetics, treatment and new insights into osteoclast function. Nat Rev Endocrinol 9, 522–536.

Soe, K., Hobolt-Pedersen, A.S., and Delaisse, J.M. (2015). The elementary fusion modalities of osteoclasts. Bone 73, 181–189.

Soriano, P., Montgomery, C., Geske, R., and Bradley, A. (1991). Targeted disruption of the c-src proto-oncogene leads to osteopetrosis in mice. Cell 64, 693–702.

Stattin, E.L., Henning, P., Klar, J., McDermott, E., Stecksen-Blicks, C., Sandstrom, P.E., Kellgren, T.G., Ryden, P., Hallmans, G., Lonnerholm, T., Ameur, A., Helfrich, M.H., Coxon, F.P., Dahl, N., Wikstrom, J., and Lerner, U.H. (2017). SNX10 gene mutation leading to osteopetrosis with dysfunctional osteoclasts. Sci Rep 7, 3012, doi:3010.1038/s41598-41017-02533-41592.

Tolar, J., Teitelbaum, S.L., and Orchard, P.J. (2004). Osteopetrosis. N Engl J Med 351, 2839-2849.

Vasikaran, S., Eastell, R., Bruyere, O., Foldes, A.J., Garnero, P., Griesmacher, A., McClung, M., Morris, H.A., Silverman, S., Trenti, T., Wahl, D.A., Cooper, C., and Kanis, J.A. (2011). Markers of bone turnover for the prediction of fracture risk and monitoring of osteoporosis treatment: a need for international reference standards. Osteoporos Int. 22, 391–420. 22, 391–420.

Wu, C.C., Econs, M.J., DiMeglio, L.A., Insogna, K.L., Levine, M.A., Orchard, P.J., Miller, W.P., Petryk, A., Rush, E.T., Shoback, D.M., Ward, L.M., and Polgreen, L.E. (2017). Diagnosis and Management of Osteopetrosis: Consensus Guidelines From the Osteopetrosis Working Group. J Clin Endocrinol Metab 102, 3111–3123.

Ye, L., Morse, L.R., Zhang, L., Sasaki, H., Mills, J.C., Odgren, P.R., Sibbel, G., Stanley, J.R., Wong, G., Zamarioli, A., and Battaglino, R.A. (2015). Osteopetrorickets due to Snx10 deficiency in mice results from both failed osteoclast activity and loss of gastric acid-dependent calcium absorption. PLoS Genet 11, e1005057.

Zhou, C., Wang, Y., Peng, J., Li, C., Liu, P., and Shen, X. (2017). SNX10 plays a critical role in MMP9 secretion via JNK-p38-ERK signaling pathway. J Cell Biochem 118, 4664–4671.

Zhou, C., You, Y., Shen, W., Zhu, Y.Z., Peng, J., Feng, H.T., Wang, Y., Li, D., Shao, W.W., Li, C.X., Li, W.Z., Xu, J., and Shen, X. (2016). Deficiency of sorting nexin 10 prevents bone erosion in collagen-induced mouse arthritis through promoting NFATc1 degradation. Ann Rheum Dis 75, 1211–1218.

Zhu, C.H., Morse, L.R., and Battaglino, R.A. (2012). SNX10 is required for osteoclast formation and resorption activity. J Cell Biochem 113, 1608–1615.

