## Supplemental Figures for "R51Q SNX10 induces osteopetrosis by promoting uncontrolled fusion of monocytes to form giant, non-functional osteoclasts"

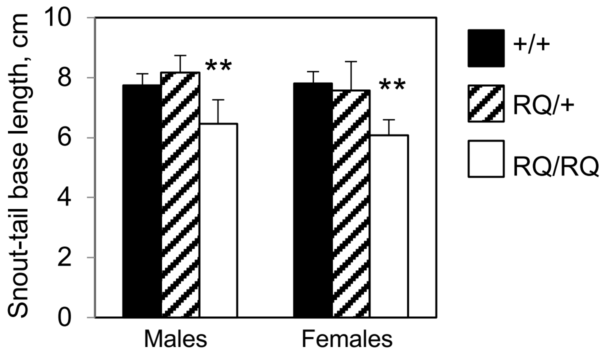


**Figure 1- figure supplement 1:** Snout-tail base length of wild-type (+/+), heterozygous (RQ/+) and homozygous (RQ/RQ) mice at age 4 weeks. Data are mean±SD, N=6-16 mice/bar. **: P≤ 0.001 by one-way ANOVA with Tukey's multiple comparisons test.

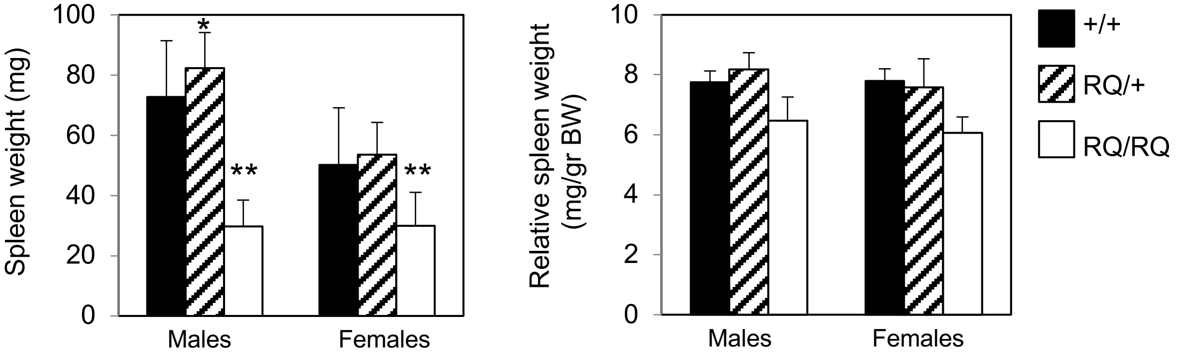


**Figure 1 – figure supplement 2:** Left: Absolute weight of spleens of mice aged 4 weeks. Right: Spleen weight normalized to body weight at 4 weeks. Data are mean±SD, N=6-16 mice/bar. *: P=0.0358 vs. +/+. **: P ≤0.0047. by one-way ANOVA with Tukey's multiple comparisons test. Data are from both lines 43 and 87.

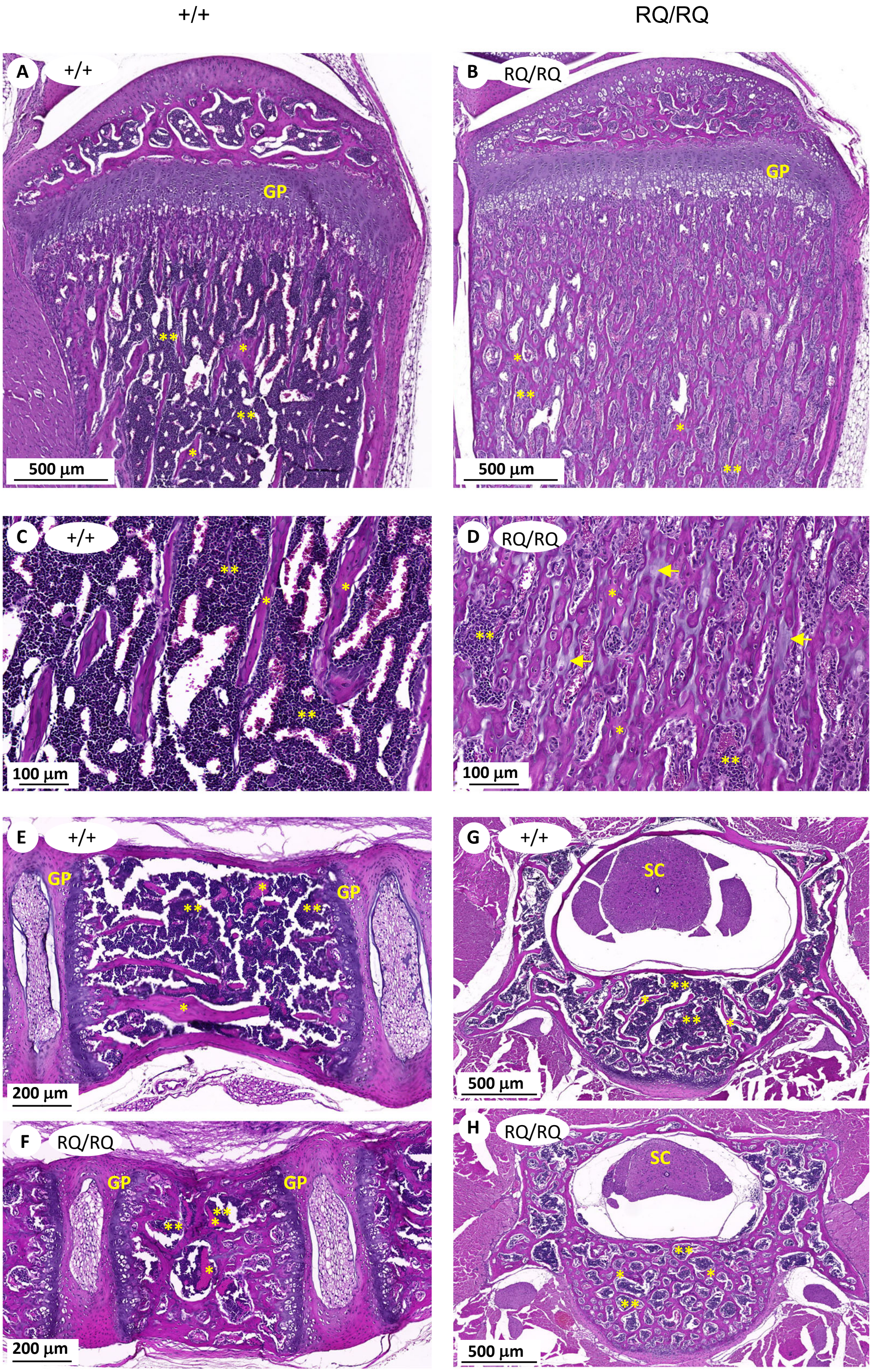


**Figure 3 – figure supplement 1: Comparison of bone structure in +/+ (Panels A, C, E, G) and RQ/RQ mice (Panels B, D, F, H) mice**. (**A, B**) Tibial growth plate region. (**C, D**) Higher magnification of the bone marrow cavity. Arrows denote residual cartilage in D. (**E, F**) Lateral sections of spinal cord vertebrae. (**G, H**) Transverse sections of vertebrae. GP; Growth plate. *: Trabecular bone. **: Bone marrow cells. SC: spinal cord. Images are representative of 4 +/+ and 4 RQ/RQ mice.

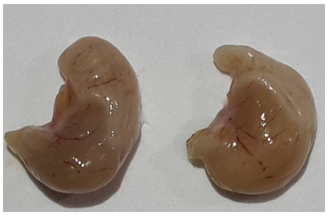


**Figure 3 – figure supplement 2: Stomachs of RQ/RQ mice display normal morphology.** Image is representative of 13 +/+ and 8 RQ/RQ mice from line 43.

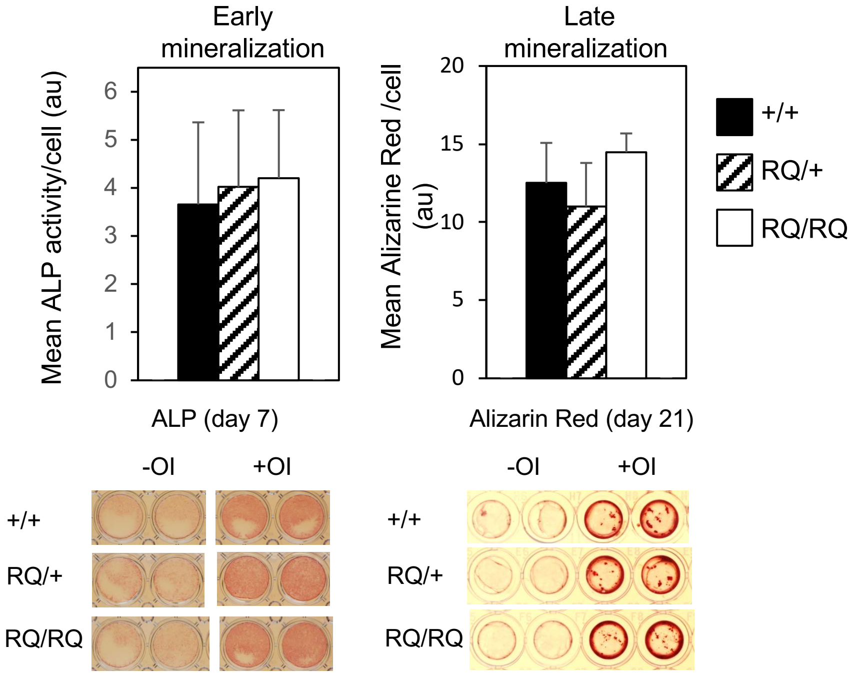


**Figure 4 – figure supplement 1: *In vitro* differentiation of calvarial mesenchymal cells to osteoblasts is unaltered by the SNX10 R51Q mutation.** Osteoblast precursors were prepared from calvarias isolated from 5-day old pups. Left: Cells were induced with ascorbic acid and β-glycerophosphate upon 80% confluency (+OI) or kept in an uninduced state (-OI). After 7 days early mineralization was quantified by alkaline phosphatase (ALP) staining. Shown are mean ±SD of 7-9 pups/bar. Right: After 21 days late mineralization and the formation of mineralized nodules was measured by alizarin red staining. Shown are mean±SD of 5-6 pups/ bar.

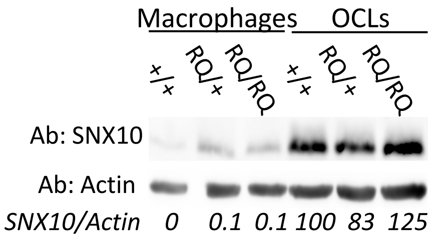


**Figure 5 – figure supplement 1:** RQ/RQ OCLs express normal amounts of SNX10 protein. +/+, RQ/+ and RQ/RQ OCLs, grown in the presence of M-CSF (macrophages) or M-CSF and RANKL (osteoclasts) were analyzed by protein blotting for SNX10 expression, with actin as a loading control. Data shown are from line 43; similar results were obtained also from line 87.

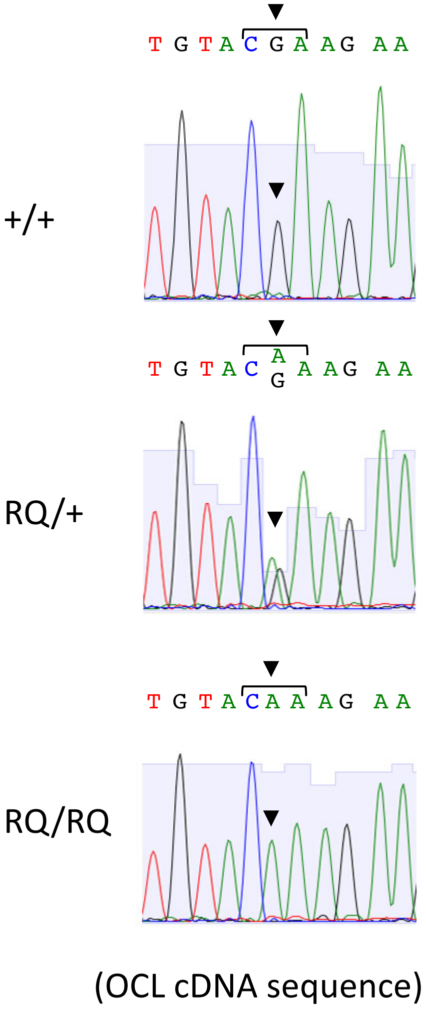


**Figure 5 – figure supplement 2: The *RQ* allele expresses mutant mRNA.** RNA from OCLs of the indicated genotypes was reverse-transcribed, amplified by PCR, and sequenced. WT and RQ/RQ samples exclusively express either WT or mutant mRNAs, respectively, while the heterozygous (RQ/+) sample expresses both. Sequence shown is in the sense orientation. The mutated triplet is marked.

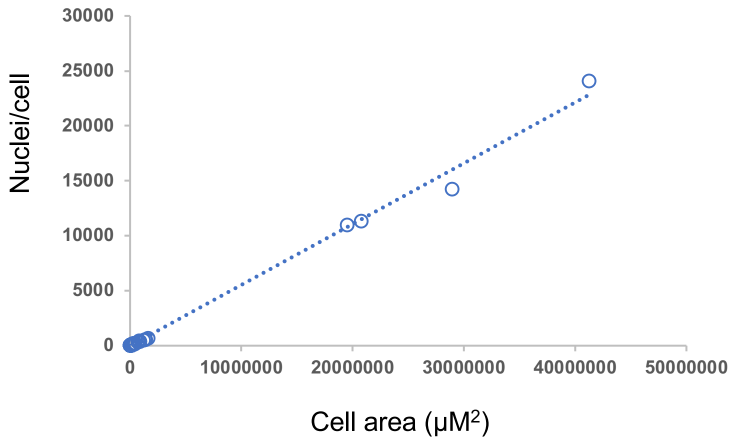


**Figure 5 – figure supplement 3: The nuclear density of giant RQ/RQ OCLs is similar to that of smaller OCLs.** Scatter diagram showing the number of nuclei and area per cell in RQ/RQ OCLs of all size ranges. N=219 RQ/RQ OCLs, $R^2$=0.9951. The large group of relatively small cells at lower left is shown also in Figure 5D. Cells were grown on plastic.
